# PPARβ/δ recruits NCOR and regulates transcription reinitiation

**DOI:** 10.1101/525576

**Authors:** Nathalie Legrand, Clemens L. Bretscher, Svenja Zielke, Bernhard Wilke, Michael Daude, Barbara Fritz, Wibke E. Diederich, Till Adhikary

## Abstract

In the absence of ligands, the nuclear receptor PPARβ/δ recruits the NCOR and SMRT corepressors, which form complexes with HDAC3, to canonical target genes. Agonistic ligands cause dissociation of corepressors and enable enhanced transcription. *Vice versa*, synthetic inverse agonists augment corepressor recruitment and repression. Both basal repression of the target gene *ANGPTL4* and reinforced repression elicited by inverse agonists are partially insensitive to HDAC inhibition. This raises the question of how PPARβ/δ represses transcription mechanistically. We show that the PPARβ/δ inverse agonist PT-S264 impairs transcription initiation by decreasing recruitment of activating Mediator subunits, RNA polymerase II, and TFIIB, but not of TFIIA, to the *ANGPTL4* promoter. Mass spectrometry identifies NCOR as the main PT-S264–dependent interactor of PPARβ/δ. Reconstitution of knockout cells with PPARβ/δ mutants deficient in basal repression results in diminished recruitment of NCOR, SMRT, and HDAC3 to PPAR target genes, while occupancy by RNA polymerase II is increased. PT-S264 restores binding of NCOR, SMRT, and HDAC3 to the mutants, resulting in reduced polymerase II occupancy. Our findings corroborate deacetylase-dependent and -independent repressive functions of HDAC3-containing complexes, which act in parallel to downregulate transcription.

## INTRODUCTION

PPARβ/δ (peroxisome proliferator activated receptor β/δ) is a type II nuclear receptor which constitutively binds to DNA as an obligate heterodimer with a retinoid X receptor (RXR). Its target genes function in lipid and glucose metabolism and also in inflammation (1, 2). An important PPAR target gene is *ANGPTL4 (angiopoietin-like 4)*, a regulator of lipid metabolism, angiogenesis, wound healing, and metastasis (3, 4). In the absence of ligands, the PPARβ/δ-RXR heterodimer represses its canonical target genes (1) *via* the recruitment of corepressors (5) such as NCOR (nuclear receptor corepressor)- and SMRT (silencing mediator of retinoid and thyroid hormone receptors)-containing complexes (6, 7, 8). Both corepressor complexes harbour the catalytic subunit histone deacetylase 3 (HDAC3) (9, 10), whose activity requires binding to NCOR or SMRT (11). Several fatty acids and their derivatives act as endogenous PPARβ/δ agonists (12, 13, 14, 15). Agonistic ligands cause dissociation of corepressors from the nuclear receptor at ligand-regulated target genes, while synthetic inverse agonists recently developed in our group lead to enhanced corepressor recruitment (4, 7, 16, 17). Basal repression of *ANGPTL4* and augmented repression in the presence of inverse agonists are largely insensitive to trichostatin A (4), an inhibitor of class I and II HDACs, suggesting an HDAC-independent repression mechanism. Induction of *ANGPTL4* transcription by activating stimuli is efficiently suppressed by PPARβ/δ inverse agonists, and this coincides with decreased binding of RNA polymerase II (RNAPII) (4). Agonists alleviate basal repression, and transcription is induced synergistically with other activating stimuli (18, 19).

The preinitiation complex (PIC) is comprised of the general transcription factors (GTFs; TFIIA, TFIIB, TFIID, TFIIE, TFIIF, and TFIIH), the Mediator complex, and RNAPII (20, 21, 22, 23, 24). Its formation is a prerequisite and a rate-limiting process for RNAPII-dependent transcription. After promoter clearance by the polymerase, additional rounds of transcription are initiated from the scaffold complex, which contains a subset of general transcription factors that remain bound to the promoter. It was shown that reinitiation of transcription from an immobilized template requires reincorporation of TFIIB, TFIIF, and RNAPII into the scaffold (25), yielding a reinitiation complex (RIC). The re-use of remaining promoter-bound GTFs supersedes the need for recurrent PIC formation and thus enables high-level transcription. Moreover, dephosphorylation of the carboxyterminal domain (CTD) of the large subunit of RNAPII after termination allows for RNAPII recycling, which is enhanced by proximity of the transcription start site (TSS) and the terminator (26). TFIIB is necessary for the formation of these gene loops (27, 28, 29). *In vitro*, human Mediator facilitates TFIIB and RNAPII recruitment (30), and its kinase module regulates reinitiation (31). Little is known about the regulation of transcripton reinitiation *in vivo* (32).

In the present study, we investigated the mechanism of transcriptional repression by PPARβ/δ inverse agonists in human cell lines. Our data show that inverse agonists specifically interfere with stimulus-dependent recruitment of TFIIB, RNAPII, and activating Mediator subunits to the *ANGPTL4* promoter, while binding of the scaffold GTFs TFIIA and TFIIH is unchanged. This suggests an impairment of the Mediator-TFIIB recruitment step, affecting RNAPII binding and reinitiation. The binding pattern of RNAPII at the *ANGPTL4* locus in the presence of an inverse agonist is similar to the pattern elicited by the transcription initiation inhibitor triptolide. We identify NCOR as the main ligand-dependent interactor of PPARβ/δ in the presence of the inverse agonist PT-S264. Strikingly, PT-S264–dependent repression is partially insensitive to both trichostatin A, a non-selective HDAC inhibitor, and also to apicidin, an HDAC3-selective inhibitor. Expression of PPARβ/δ mutants in PPARβ/δ knockout cells identified amino acid residues required for basal repression. These PPARβ/δ mutants revealed diminished NCOR, SMRT, and HDAC3 binding to chromatin in the basal state, concomitant with increased RNAPII binding. Recruitment of NCOR, SMRT, and HDAC3 was restored by the inverse agonist, as was RNAPII loss and repression of transcription. Repression of PPAR target genes by these mutant receptors was largely insensitive to HDAC inhibition. Our data show that chromatin-bound NCOR and SMRT complexes downregulate transcription reinitiation, and possibly initiation, *via* both deacetylase-dependent and -independent mechanisms.

## MATERIALS AND METHODS

### Antibodies

The following antibodies were used in this study: HDAC3, Santa Cruz no. sc-11417, rabbit polyclonal, ChIP, RRID AB_2118706; LDH, Santa Cruz no. sc-33781, rabbit polyclonal, immunoblot, RRID AB 2134947; MED1, Santa Cruz no. sc-8998, rabbit polyclonal, ChIP, RRID AB_2144021; MED13L, Bethyl no. A302-420A, rabbit polyclonal, ChIP, RRID AB 1907303; MED26, Santa Cruz no. sc-48776, rabbit polyclonal, ChIP, RRID AB 782277; NCOR, Abcam no. ab24552, rabbit polyclonal, ChIP, RRID AB 2149005; NCOR, Bethyl no. A301-145A, rabbit polyclonal, ChIP, RRID AB_873085; NCOR, Thermo no. PA1-844A, rabbit polyclonal, immunoblot, RRID AB 2149004; IgG fraction, Sigma no. I5006, rabbit polyclonal, ChIP, RRID AB 1163659; PPARα, Santa Cruz no. sc-9000, rabbit polyclonal, ChIP and immunoblot, RRID AB 2165737; PPARβ/δ, Santa Cruz no. sc-7197, rabbit polyclonal, ChIP, RRID AB_2268420; PPARβ/δ, Santa Cruz no. sc-74517, mouse monoclonal, immunoblot, RRID AB 1128604; PPARγ, Santa Cruz no. sc-7196, rabbit polyclonal, ChIP and immunoblot, RRID AB 654710; RPB1 (RNAPII large subunit) CTD (33), Biolegend no. 8WG16, mouse monoclonal, ChIP, RRID AB 2565554; RPB1 unphosphorylated CTD (34), Ascenion no. 1C7, rat monoclonal, ChIP, RRID AB 2631402; RPB1 Ser5-phosphorylated CTD (35), Ascenion no. 3E8, rat monoclonal, ChIP, RRID AB 2631404; RPB1 Ser2-phosphorylated CTD (35), Ascenion no. 3E10, rat monoclonal, ChIP, RRID AB 2631403; RPB1 NTD, Santa Cruz no. sc-899, rabbit polyclonal, ChIP, RRID AB 632359; RPB1 NTD, Santa Cruz no. sc-9001, rabbit polyclonal, ChIP, RRID AB 2268548; RXR, Santa Cruz no. sc-774, rabbit polyclonal, ChIP, RRID AB 2270041; SMRT, Abcam no. ab24551, rabbit polyclonal, ChIP, RRID AB_2149134; TBLR1, Novus no. NB600-270, rabbit polyclonal, ChIP, RRID AB 10001343; TBP, Santa Cruz no. sc-273, rabbit polyclonal, ChIP, RRID AB 2200059; TFIIA, Santa Cruz no. sc-25365, rabbit polyclonal, ChIP, RRID AB 2116529; TFIIB, Santa Cruz no. sc-225, rabbit polyclonal, ChIP, RRID AB 2114380; TFIIH, Santa Cruz no. sc-293, rabbit polyclonal, ChIP, RRID AB_2262177.

### Compounds

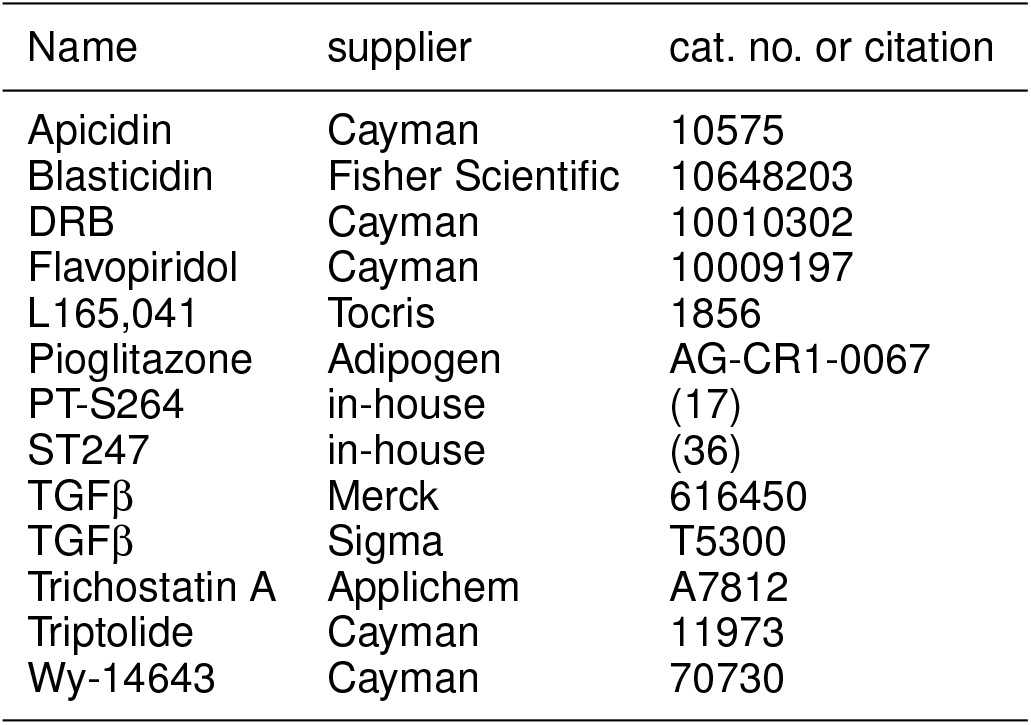

### Cell culture and treatment

φNX cells (37) were cultivated in DMEM with 10% FBS. MDA-MB231-*luc2* and Caki-1 cells were cultivated in McCoy’s 5A medium with 10% FBS. The compound or compounds were added for 6 h (expression analyses) or 30 min (ChIP assays) to the cultures. Control populations were supplied with an equivalent volume of solvent (DMSO for low molecular weight compounds, PBS with 0.1 % (w/v) fatty acid free BSA for TGFβ).

### Genetic deletion and stable transfection

Vectors for parallel expression of a guide RNA and the Cas9 nuclease were obtained from Santa Cruz Biotechnology. The sequences targeting the *PPARD* coding region were TCGTACGATCCGCATGAAGC (1), CCCTGTGCAGCTATCCGTTT (2), and AACACTCACCGCCGTGTGGC (3). MDA-MB231-*luc2* cells were transfected using Lipofectamine 2000 (Life Technologies) according to the manufacturer’s instructions with the guide/Cas9 expression vectors together with pMSCVbsd (38) for selection of transfected cells. After four hours, the cells were supplemented with fresh medium, and blasticidin was added after 30 hours at a concentration of 10 μg/ml. After ten days, single cells were seeded using limiting dilution in cell-free medium conditioned by the parental cell line. Wells with more than one colony were terminated. Clones were expanded and screened *via* RT-qPCR (loss of *ANGPTL4* repression by PT-S264) and immunoblotting against PPARβ/δ. The 2B3 clone was transfected with pWZLneo-ecoR (39). After 24 hours, the cells were selected with 500 μg/ml G418 for 14 days.

### *PPARD* expression vectors

The retroviral vector pMSCVbsd (38) was used to clone the *PPARD* cDNA into the *Bgl*II and *Xho*I sites. Terminal deletions were introduced *via* PCR with specific primers, and the fragment was re-inserted into the empty pMSCVbsd vector. Alterations in the cDNA sequence were introduced with site-directed mutagenesis. All plasmid preparations used for retroviral transduction were validated by sequencing of the entire cDNA inserts. Subsequent new plasmid preparations were validated by resequencing. Vectors are available from Addgene.

### Retroviral transduction

For production of ecotropic retroviruses, pMSCVbsd-*PPARD* vectors or the empty vector were transfected into subconfluent φNX-eco packaging cells (37). After four hours, 7 ml of fresh medium were added. Twenty-four hours later, the supernatant was harvested, centrifuged at 800×g, and 3 ml aliquots were snap frozen. The φNX-eco cells received fresh medium, and a second supernatant was harvested 24 hours later. Freshly seeded MDA-MB231-*luc2*-2B3 cells ectopically expressing the ecotropic receptor were incubated with 2 ml of fresh medium, 3 ml of φNX supernatant, and 4 μg/ml polybrene for 24 hours. Selection was performed with 10 μg/ml blasticidin for ten days, and cells were subsequently cultured in the presence of blasticidin.

### RNA isolation, cDNA synthesis, and quantitative RT-PCR

Total RNA was isolated with the NucleoSpin RNA kit (Macherey&Nagel, no. 740955) according to the manufacturer’s instructions; the DNase digestion and desalting steps were omitted. Complementary DNA synthesis was carried out with the iScript cDNA Synthesis Kit (Bio-Rad, no. 170-8891SP) according to the manufacturer’s instructions with 500 ng of purified RNA per sample in 20 μl. Quantitative PCR analyses were performed in three technical replicates per sample using ABsolute SYBR Green master mix (Thermo Scientific, no. AB-1158B) in a total reaction volume of 10 μl in Mx3000p and Mx3005 thermocyclers (Agilent). Prior to PCR, cDNA samples were diluted 1:10, and 4.75 μl were used per reaction. The *RPL27* transcript was used for normalization. RT-qPCR was carried out with primer concentrations of 250 nM each. The primer sequences are available in the supplementary information file (supp. table S1). Ct values were normalised to the *RPL27* transcript; to retain information about *ANGPTL4* expression levels, the mean *RPL27* Ct value of all samples in the respective assay was added back where suitable.

### Chromatin immunoprecipitation (ChIP)-qPCR

ChIP was essentially performed as described previously (40, 41). Fixation was performed with 1 % formaldehyde in media for 10 min at room temperature followed by quenching with 125 mM glycin for 5 min. Cells were lysed in buffer L1 (5 mM PIPES pH 8.0, 85 mM KCl, 0.5% (v/v) NP40, protease inhibitor mix (Sigma, no. P8340, 1:1000) for 20–40 min on ice. Nuclei were resuspended in ChIP RIPA buffer (10 mM Tris-HCl pH 7.5, 150 mM NaCl, 1 % (v/v) NP40, 1 mM EDTA) supplemented with 1:1000 protease inhibitor mix (Sigma), incubated on ice for 10–20 min and sheared with a Branson S250D Sonifier (Branson Ultrasonics) using a microtip in 1 ml aliquots in 15 ml conical tubes. 40–52 pulses of 1 s, 4 s pause, 20% amplitude were applied with cooling of the sample in an ice-ethanol mixture or in a 15 ml tube cooler (Active Motif, no. 53077). A 15 min 17,000×g supernatant was precleared with 10 μg of IgG coupled to 100 μl of blocked sepharose slurry (see below) for 45 min at 4 °C with agitation. IP was carried out with 300 μl of precleared chromatin, equivalent to 6–8×10^6^ cells. ChIP was performed using 4 μg of antibody per sample. For precipitation, a mixture of protein A and protein G sepharose (GE Healthcare life sciences, no. 1752800 and no. 1706180) or pure protein A sepharose (Zymed) was washed twice with ChIP RIPA buffer and blocked with 1 g/l BSA and 0.4 g/l sonicated salmon sperm DNA (Life Technologies no. 15632011) overnight. 50 μl of blocked bead slurry (1:1 volume ratio with liquid phase) were used per IP. Samples were washed once in buffer I (20 mM Tris pH 8.1; 150 mM NaCl; 1 % (v/v) Triton X-100; 0.1 % (w/v) SDS; 2 mM EDTA), once in buffer II (20 mM Tris pH 8.1; 500 mM NaCl; 1 % (v/v) Triton X-100; 0.1 % (w/v) SDS; 2 mM EDTA), twice in buffer III (10 mM Tris pH 8.1; 250 mM LiCl; 1 % (v/v) NP40; 1 % (w/v) sodium deoxycholate; 1 mM EDTA) on ice and twice in Qiagen buffer EB (10 mM Tris pH 8.0; no. 19086) at room temperature. Immune complexes were eluted twice with 100 mM NaHCO3 and 1 % (w/v) SDS under agitation. Eluates were incubated overnight at 65 °C after adding 10 μg of RNase A and 20 μg of proteinase K in the presence of 180 mM NaCl, 35 mM Tris-HCl pH 6.8 and 9 mM EDTA. An input sample representing 1 % of the chromatin used per IP was reverted in parallel. Samples were purified using the Qiagen PCR purification kit according to the manufacturer’s instructions. ChIP-qPCR was performed in three technical replicates per sample with the ABsolute SYBR Green master mix (Thermo Scientific, no. AB-1158B) in in Mx3000p and Mx3005 thermocyclers (Agilent). Quantitative PCR was carried out with primer concentrations of 250 nM each. The primer sequences are available in the supplementary information file (supp. table S2).

### ChIP–mass spectrometry (ChIP-MS)

Two technical replicates each were performed by Active Motif Epigenetic Services according to a published protocol (42) from the following MDA-MB231-*luc2* samples: IgG, PT-S264 treatment (0.3 μM); α-PPARβ/δ, PT-S264 treatment (0.3 μM); α-PPARβ/δ, L165,041 treatment (1.0 μM). The unspecific IgG pool from rabbit was obtained from Sigma-Aldrich (no. I5006), the PPARβ/δ antibody from Santa Cruz Biotechnology (sc-7197, rabbit polyclonal). Candidate proteins were filtered with scaffold Viewer 4.8.3 (Proteome Software) according to the following criteria: At least two unique peptides in total and a maximum of two unique peptides each in the negative control replicates (IgG). The protein and the peptide thresholds were each set to 80 %.

## RESULTS

### PPARβ/δ inverse agonists interfere with formation of an initiation complex

PT-S264 is an inverse PPARβ/δ agonist with improved repressive properties and stability (17). We first tested whether it affects initiation complex formation like the previously used inverse agonist ST247 (36), which reduces RNAPII binding to the *ANGPTL4* promoter (4). TGFβ is a strong inducer of *ANGPTL4* transcription in human cells (18). Treatment with PT-S264 strongly reduced RNAPII (large subunit RPB1) binding to the *ANGPTL4* TSS as shown by scanning ChIP-qPCR using closely spaced primer pairs (fig. 1A) both in the presence and in the absence of TGFβ in Caki-1 cells. We used the 8WG16 antibody, which in our hands detects total RNAPII with superior specificity to other RNAPII antibodies (supp. fig. S1). The binding pattern of RNAPII in PT-S264–treated cells is similar to the binding pattern observed in cells treated with the transcription initiation inhibitor triptolide, which however is more effective in reducing RNAPII binding compared to PT-S264 (upper panels of fig. 1A). In contrast, treatment of the cells with the CDK9 inhibitors DRB and flavopiridol, which interfere with elongation, does not prevent TGFβ-induced RNAPII accumulation at the TSS (blue and green lines in the lower panels of fig. 1A). This indicates that PT-S264 prevents RNAPII recruitment to the *ANGPTL4* promoter in a manner similar to the effect of triptolide. Taken together, these observations show that PT-S264 impinges on transcription initiation of *ANGPTL4*. Notably, ChIP-qPCR data obtained with antibodies against the RPB1 CTD phosphorylated at serine 5 (initiating RNAPII) or serine 2 (elongating RNAPII) do not support an effect of PT-S264 on serine 2 phosphorylation when taking total RNAPII enrichment into account (supp. fig. S1), which again implies that the inverse agonist diminishes RNAPII recruitment and furthermore argues against an effect of PT-S264 on RPB1 CTD phosphorylation. We therefore conclude that the formation of an initiation complex is affected by PT-S264.

**Figure 1.**
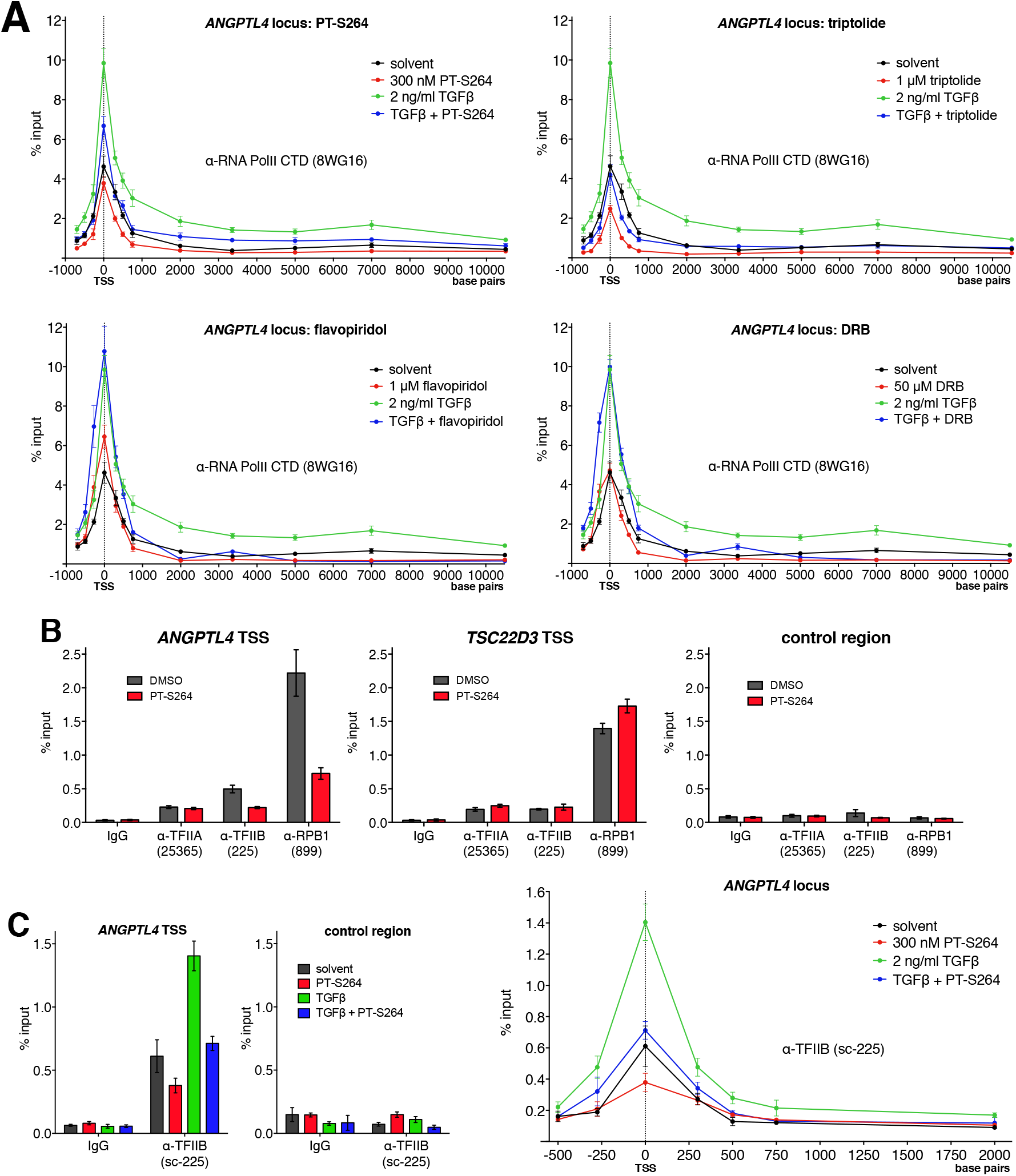
PPARβ/δ inverse agonists suppress transcription initiation of *ANGPTL4* and recruitment of TFIIB to a scaffold complex. Caki-1 cells were treated for 30 min as indicated and subjected to ChIP-qPCR analysis. **A**: Scanning ChIP-qPCR of the *ANGPTL4* locus. An antibody against the CTD of the large subunit of RNAPII was used. **B**: Binding of TFIIA, TFIIB, and RPB1 at the *ANGPTL4* and *TSC22D3* TSSs. **C**: TFIIB binding at the *ANGPTL4* TSS in the presence of TGFβ, PT-S264, or both. The right panel shows scanning ChIP around the *ANGPTL4* TSS from the same samples. Commercially available antibodies are denominated by the terms in parentheses. Means and standard deviations of three technical replicates from representative ChIP assays are plotted.

In order to clarify how PT-S264 interferes with RNAPII recruitment, we performed further experiments with the Caki-1 line. These cells highly express *ANGPTL4* (4) and hence allow for ChIP-based detection of GTFs at the endogenous locus. To address which step of initiation complex formation is affected, we measured the occupancy of GTFs at the *ANGPTL4* TSS in the presence or in the absence of PT-S264. Binding of TFIIB and RNAPII is reduced upon treatment with PT-S264 compared to the control. However, the occupancy of TFIIA is not altered (fig. 1B), suggesting that the repressive mechanism affects the transition from the scaffold complex to the RIC. PIC formation may be impaired too, which cannot be inferred from these data obtained after short-term treatment. The effect of PT-S264 is PPAR-dependent, as TFIIB and RNAPII binding remain unchanged at the TSS of *TSC22D3*, which is not a PPAR target. In the presence of TGFβ, TFIIB binding to the *ANGPTL4* TSS is increased, and this is counteracted by PT-S264 (fig. 1C).

The Mediator complex facilitates the rate-limiting step of TFIIB recruitment, and both Mediator and TFIIB are necessary for RNAPII binding in a reconstituted human transcription system (30) as well as in *S.cerevisiae in vivo* (43). Initial ChIP-qPCR experiments after treatment with the previously used inverse agonist ST247 show that both TFIIH, another GTF present in the scaffold, and MED1 (Mediator subunit 1) bind to the *ANGPTL4* TSS after TGFβ treatment. However, in the presence of the inverse agonist ST247, MED1 recruitment is blocked (supp. fig. S2); TBP ChIP data were not conclusive and show an enrichment pattern similar to MED1. The TFIIA and TFIIH ChIPs show that inverse agonists do not generally block recruitment of PIC components. This led to the hypothesis that PPARβ/δ inverse agonists affect Mediator binding to the promoter. Scanning ChIP-qPCR analyses of Caki-1 cells treated with PT-S264, TGFβ, or both show that PT-S264 counteracts TGFβ-stimulated recruitment of MED1 (fig. 2A) and MED26 (fig. 2B) around the *ANGPTL4* TSS. Both MED1 and MED26 are enriched in forms of Mediator that are permissive for transcription (44, 45, 46) Since the Mediator kinase module is detected at promoters in metazoans (47), we performed additional scanning ChIP across the *ANGPTL4* TSS with an antibody against MED13L, a Mediator subunit that resides in the kinase module in the presence of MED26 (48). Recruitment of MED13L upon stimulation with TGFβ but not after simultaneous treatment with the inverse agonist PT-S264 (fig. 2D) is detected in the same manner as MED1 (fig. 2A and supp. fig. S2), MED26 (fig. 2B), TFIIB (fig. 1B), and RNAPII (fig. 1A and supp. fig. S1). Taken together, our data suggest that MED1, MED13L, and MED26 binding occur in a transcriptionally permissive state induced by TGFβ, which is prevented in the presence of PT-S264. We conclude that PPARβ/δ inverse agonists perturb transcription initiation at the *ANGPTL4* TSS by interfering with TFIIB–RNAPII recruitment to promoter-bound GTFs, and this is achieved by preventing recruitment of MED1, MED13L, and MED26 to the promoter.

**Figure 2.**
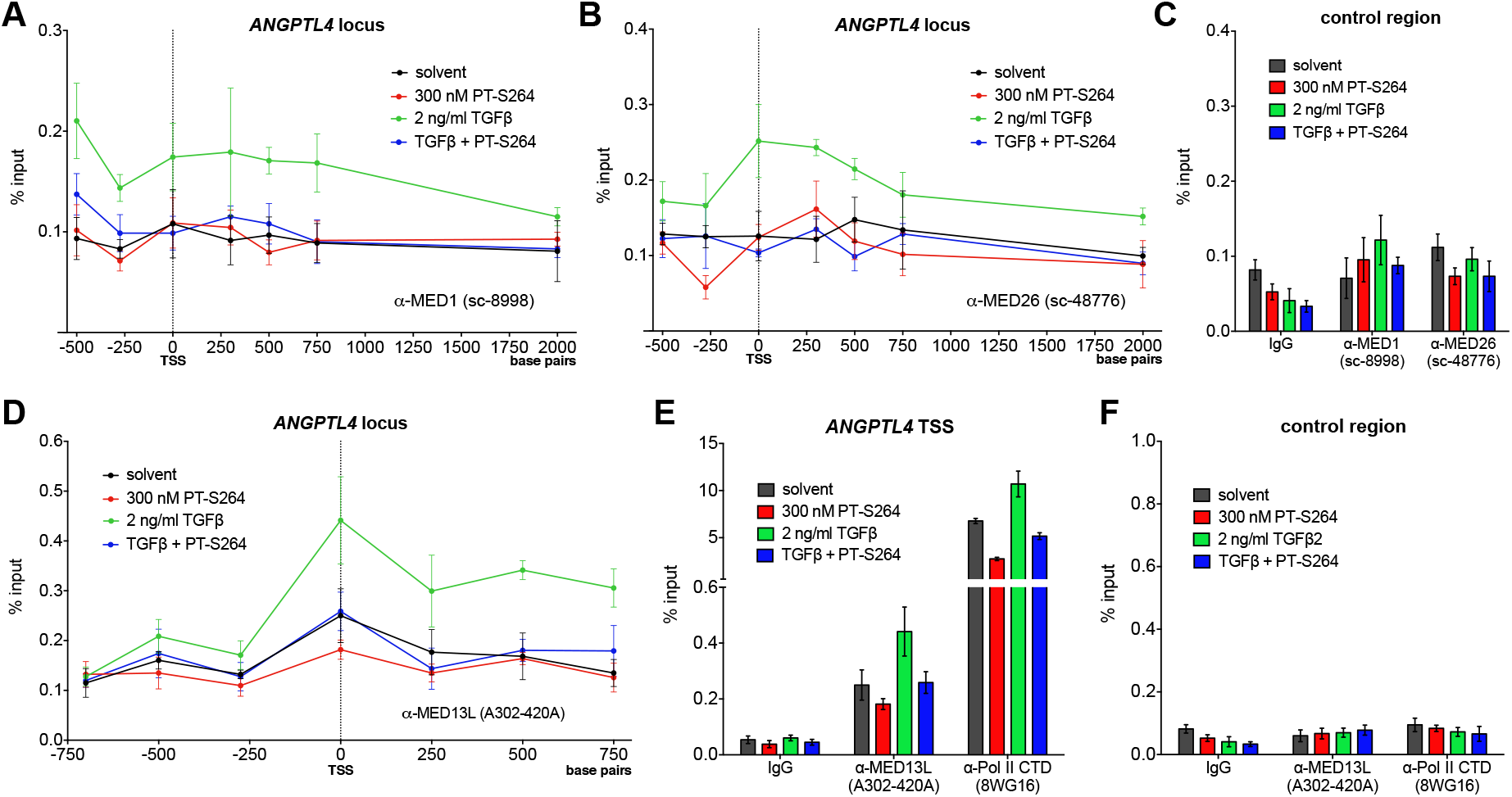
PPARβ/δ inverse agonists prevent TGFβ-stimulated recruitment of Mediator subunits to the *ANGPTL4* promoter. Caki-1 cells were treated for 30 min as indicated and subjected to scanning ChIP-qPCR analysis with primer pairs that amplify regions close to the TSS of *ANGPTL4*. Antibodies against MED1(**A**), MED26 (**B**), and MED13L (**D**) were used. Panels **C** and **F** show ChIP-qPCR data from these samples using a negative control region primer pair. In **E**, RNAPII binding is shown as an additional control. Commercially available antibodies are denominated by the terms in parentheses. Means and standard deviations of three technical replicates from representative ChIP assays are plotted.

### PT-S264 augments NCOR binding to PPARβ/δ

We previously found that depletion of NCOR by RNAi alleviates basal repression to the same extent as PPARβ/δ depletion but does not prevent repression by an inverse agonist (4). However, RNAi-mediated partial depletion may not suffice for full loss of function, especially if the affinity of PPARβ/δ binding to NCOR is considerably high in the presence of an inverse agonist. To identify candidate repressors in an unbiased approach, ChIP-mass spectrometry (ChIP-MS) according to the RIME (rapid immunoprecipitation and mass spectrometry of endogenous proteins) protocol (42) was performed with an antibody against PPARβ/δ. We used chromatin from MDA-MB231-*luc2* cells, in which PPARβ/δ target gene repression by inverse agonists is particularly strong (supp. fig. S2 and (4)). PPARβ/δ and RXR were robustly identified both in the presence of the inverse agonist PT-S264 and the agonist L165,041 (see table 1). Importantly, NCOR (encoded by the *NCOR1* gene) and SMRT (encoded by the *NCOR2* gene) were identified as interactors only in the presence of PT-S264 but not in the presence of L165,041. Other known transcriptional repressors were not detected. The NCOR2 protein cluster revealed 15 and 7 unique peptides for NCOR after treatment with PT-S264 in the two technical replicates, while only 4 and 3 peptides were identified for SMRT. This finding may indicate that NCOR binding to PPARβ/δ is dominant over SMRT binding in MDA-MB231-*luc2* cells in the presence of PT-S264. Of note, TBL1XR1, a subunit of NCOR and SMRT complexes (9), was also only detected in the presence of PT-S264 but not in the presence of L165,041. Other NCOR and SMRT complex subunits such as HDAC3 were not detected. This could be due to insufficient sensitivity, the destructive nature of chromatin sample preparation, or both.

**Table 1.**
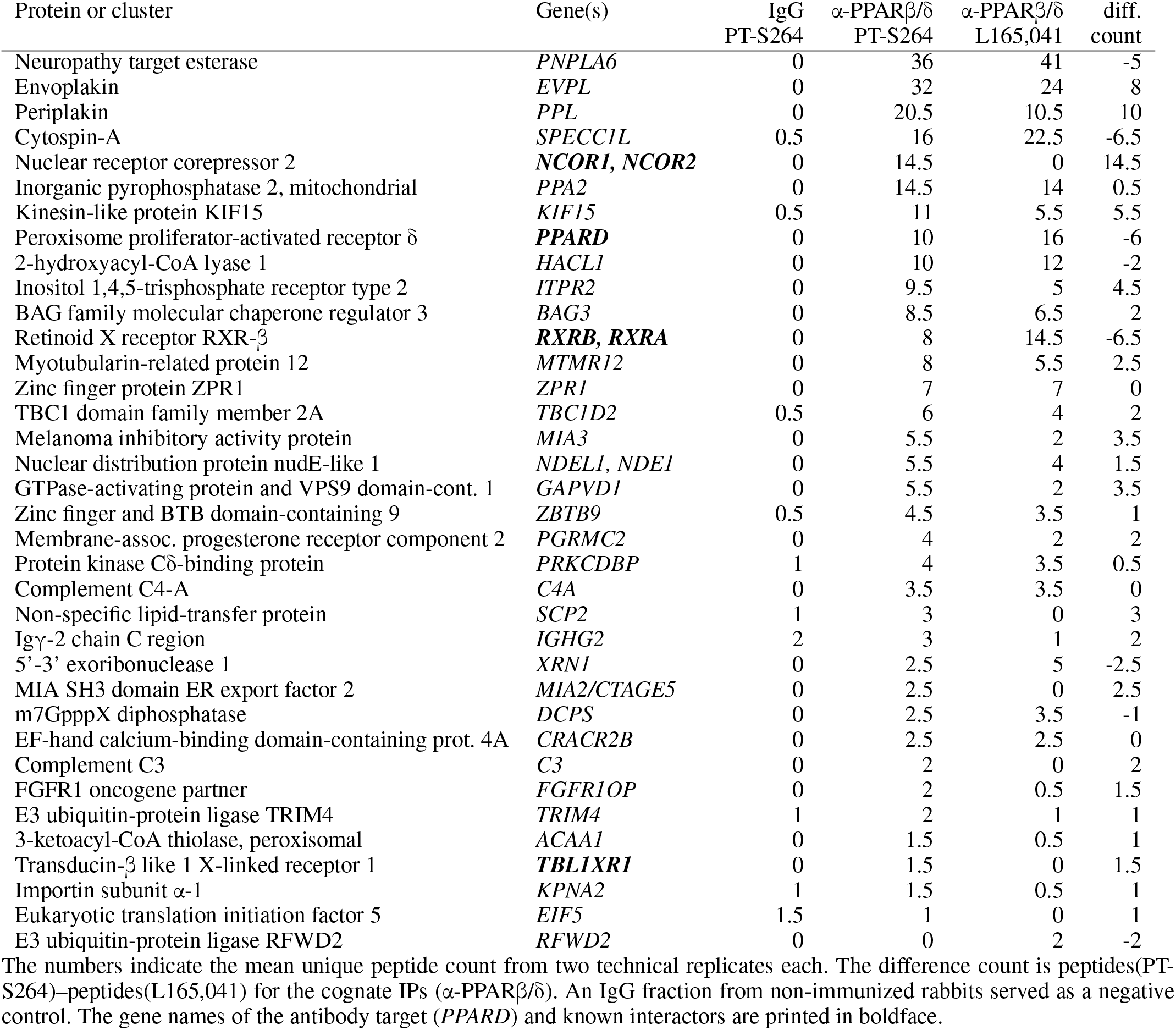
PPARβ/δ interactors in the presence of the inverse agonist PT-S264 or the agonist L165,041 (RIME ChIP-MS analysis).

Next, we measured NCOR binding to the PPAR β/δ binding sites (PPREs) of the target genes *ANGPTL4* and *PDK4* (1) in the presence and in the absence of PT-S264 by ChIP-qPCR using two different antibodies. In agreement with ChIP-MS data, NCOR is present at the *ANGPTL4* PPREs. Interestingly, we find increased binding of NCOR after treatment with the inverse agonist PT-S264 both in the presence and in the absence of TGFβ (fig. 3A). Further ChIP-qPCR analyses of several NCOR and SMRT complex subunits are in line with the ChIP-MS results, showing increased recruitment to the PPARβ/δ binding site of *ANGPTL4* in the presence of PT-S264 and decreased recruitment in the presence of L165,041 (fig. 3B). This includes PT-S264–dependent recruitment of HDAC3, which is clearly expressed in these cells (4). In Caki-1 cells, *ANGPTL4* expression is modulated by L165,041 and PT-S264 in the same manner as in MDA-MB231-*luc2* cells; the amplitude of the effects is weaker in Caki-1, while expression in the basal state is considerably higher (supp. fig. S3). Increased NCOR recruitment to the *ANGPTL4* PPREs is also observed in Caki-1 cells upon treatment with PT-S264 (supp. fig. S4). Due to its marked contribution to basal repression (4) and its enhanced recruitment to chromatin-bound PPARβ/δ in the presence of PT-S264, we postulate that NCOR mediates downregulation of *ANGPTL4* by PPARβ/δ in the basal state and in the presence of PPARβ/δ inverse agonists. This raises the question whether the interaction of NCOR and PPARβ/δ is necessary for PT-S264–dependent repression.

**Figure 3.**
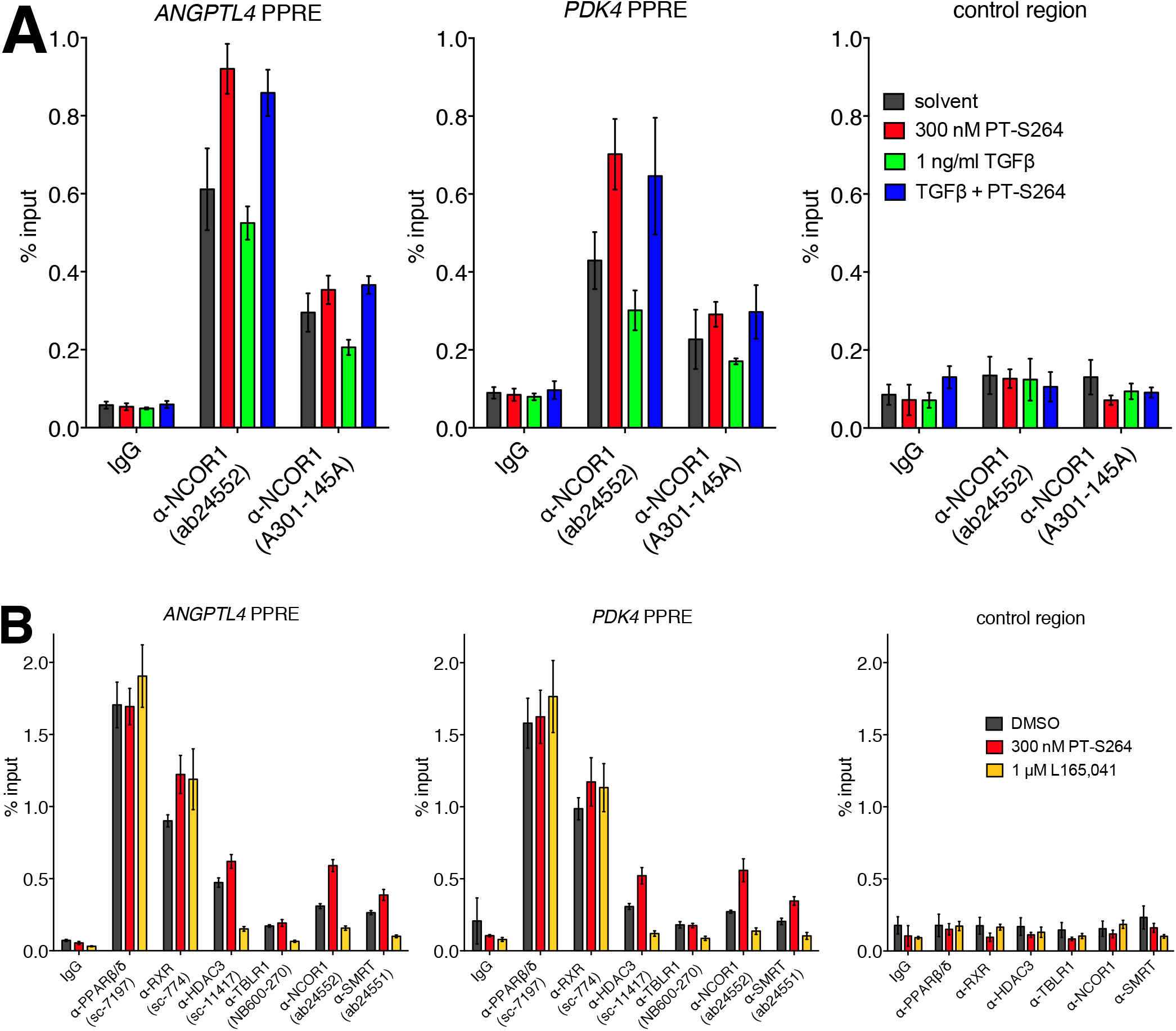
NCOR and SMRT complex subunits are recruited to PPARβ/δ in the presence of PT-S264. MDA-MB231-*luc2* cells were treated as indicated for 30 min, and ChIP-qPCR was performed with antibodies against subunits of HDAC3-containing complexes using primer pairs encompassing the *ANGPTL4* and *PDK4* PPREs (+3,500 or −12,200 bp relative to the TSS of the gene, respectively). **A**: Binding of NCOR in the presence of 300 nM PT-S264, 1 ng/ml TGFβ, or both. Two different antibodies against NCOR were used. **B**: Binding of PPARβ/δ, RXR, HDAC3, TBLR1, NCOR, and SMRT in the presence of solvent (DMSO), 300 nM PT-S264, or 1 μM L165,041. Both panels: Representative ChIP data are plotted. Commercially available antibodies are denominated by the terms in parentheses. Values denote means and standard deviations from three technical replicates.

### Generation and characterisation of PPARβ/δ knockout clones

To test whether NCOR is functionally important for PT-S264–mediated repression, we tried to disrupt NCOR expression through genetic deletion with the CRISPR-Cas9 system but were repeatedly unable to obtain NCOR knockout clones. We therefore aimed to disrupt NCOR binding to PPARβ/δ through mutations of the receptor as an alternative strategy. As a prerequisite for screening PPARβ/δ mutants, we introduced frameshift mutations to the *PPARD* coding region in MDA-MB231-*luc2* cells using the CRISPR-Cas9 system. Four different PPARβ/δ knockout clones were isolated (2B2, 2B3, 2A3, 2A6; see supp. fig. S5A). Neither PPARα nor PPARγ expression was disrupted in the four clones (supp. fig. S5B,C). ChIP-qPCR confirmed the absence of PPARβ/δ at the *ANGPTL4* and *PDK4* loci (fig. 4). Interestingly, binding of PPARα and PPARγ to the *ANGPTL4* and *PDK4* loci, each of which harbours three adjacent PPRE sequences (1, 14, 18), is increased in PPARβ/δ knockout cells, suggesting that PPARα, PPARβ/δ, and PPARγ compete for the same binding sites. However, relative to wildtype cells, binding of RXR is reduced in PPARβ/δ KO cells. This finding indicates that PPARβ/δ is the main factor that binds to the PPREs of the *ANGPTL4* and *PDK4* genes in MDA-MB231-*luc2*. Consistently, both induction of the *ANGPTL4* transcript by the synthetic agonist L165,041 and repression by the inverse agonist PT-S264 were disrupted in the knockout clones (supp. fig. S6). In contrast, the PPARγ agonist pioglitazone retained its function and stimulated *ANGPTL4* expression. Surprisingly (for unknown reasons), the PPARα agonist Wy-14643 did not induce expression of the *ANGPTL4* gene in the KO clones (supp. fig. S6).

**Figure 4.**
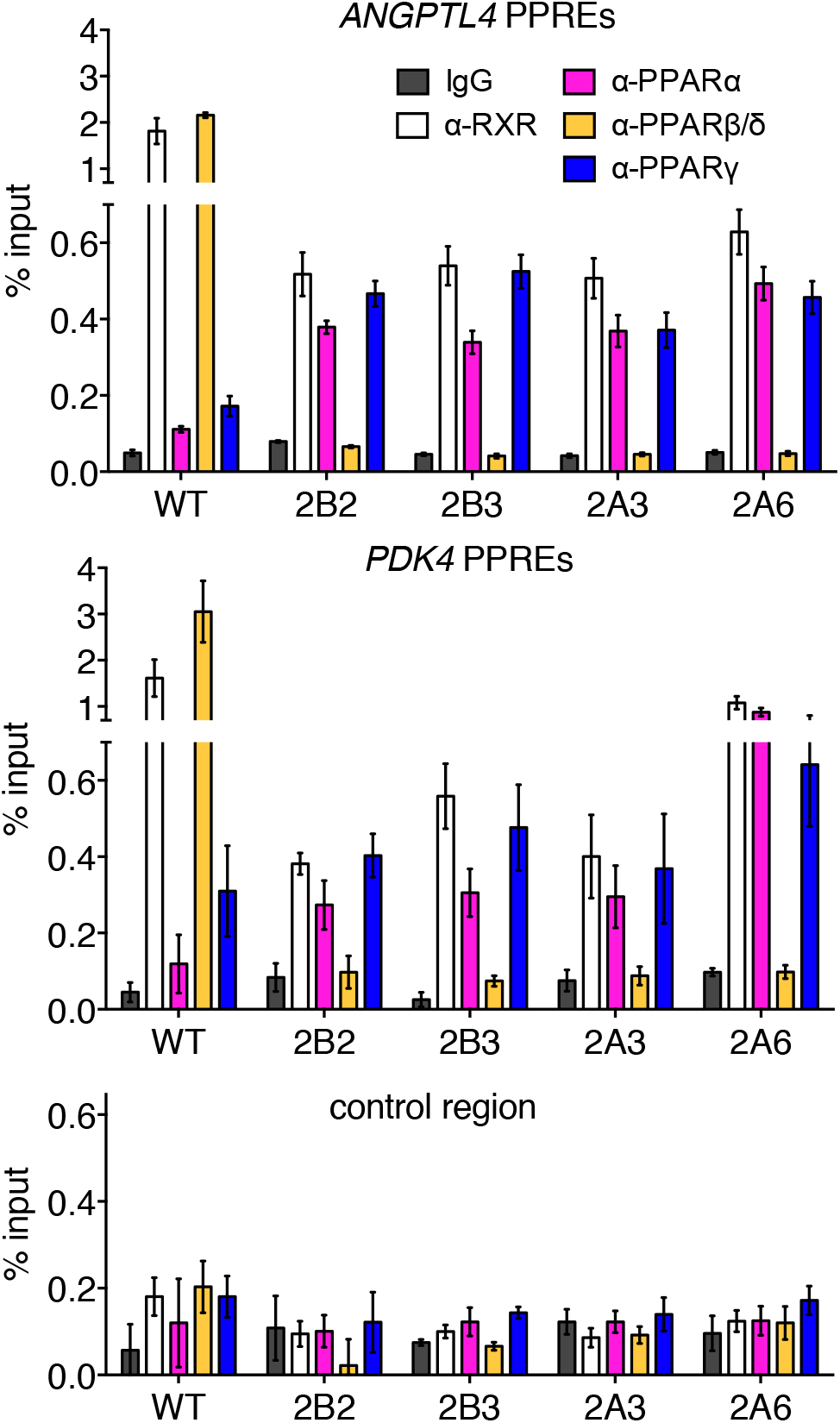
Analysis of PPAR isoform binding to chromatin in PPARβ/δ knockout clones. ChIP-qPCR analysis of PPARα, PPARβ/δ, PPARγ, and RXR binding at the *ANGPTL4* and *PDK4* loci in the four MDA-MB231-*luc2* PPARβ/δ knockout clones (2B2, 2B3, 2A3, and 2A6). Parental cells (WT) were processed in parallel. A representative experiment is shown. The error bars denote standard deviations from three technical replicates.

### Functional reconstitution of PPARβ/δ in knockout cells

In contrast to other PPARβ/δ knockout clones we obtained, the 2B3 clone neither showed growth defects, nor poor viability (observed in the 2A6 clone after several passages, data not shown), nor stable integration of the Cas9 expression vector (2B2, supp. fig. S5E) or of the transiently transfected blasticidin resistance marker (2A3; supp. fig. S5F). We therefore used the 2B3 clone for rescue experiments with expression vectors that encode for wildtype PPARβ/δ. Sequencing of cloned PCR fragments indicates that guide RNAs 1 and 3 result in single-base deletions (no. 1) or insertions (no. 3; sequences from clone 2B3 are shown in supp. fig. S5G). In line with previous observations (49), ectopic expression of PPARβ/δ driven by the strong cytomegalovirus (CMV) promoter led to formation of intracellular aggregates and failed to restore ligand function (data not shown). We stably transfected the 2B3 clone with a vector encoding the murine ecotropic retroviral receptor (39), allowing for ecotropic retroviral infection. The resulting 2B3-ecoR cells were then transduced with pMSCV vectors to express wildtype or mutant PPARβ/δ, driven by the comparably weak viral long terminal repeats. Expression of the mutants was confirmed on protein level (supp. fig. S7). We assume that PPARβ/δ is weakly expressed in all cellular models we analysed due to low signals on both RNA and protein levels. The level of PPARβ/δ ectopically expressed from pMSCV was above that of wildtype cells (supp. fig. S7). Nevertheless, both the activation and repression functions of PPARβ/δ were restored, albeit to lesser extent than that observed in wildtype cells (supp. fig. S8).

### A reconstitution screen identifies PPARβ/δ mutants deficient in ligand response and basal repression

Using PPARβ/δ knockout cells, it is possible to screen for mutants which are compromised in their repressive function. We generated a panel of 80 mutants that encode for truncated proteins or proteins with one or more substitutions of amino acid residues. We preferentially mutated residues at the surface of the LBD (ligand binding domain) since the LBD is involved in NCOR interactions (50, 51, 52), and the affinity of corepressor-derived peptides to the PPARβ/δ LBD is enhanced by inverse agonists (7). Published X-ray crystallography structural data of the PPARβ/δ LBD bound to an agonist (PDB ID `3TKM` (53)), the PPARα LBD bound to an inverse agonist (PDB ID `1KKQ` (51)), and the PPARγ-RXR heterodimer (PDB ID `3DZY` (54)) were used as reference for choosing candidate residues. The panel also includes a negative control, C91A-E92A, a double mutant of residues critical for DNA binding (55). This mutant should be defective in all functions of DNA-bound PPARβ/δ (basal repression, induction by agonist, repression by inverse agonist). We then performed a functional screen, intending to identify residues of PPARβ/δ which are necessary for basal and PT-S264–regulated repression of the *ANGPTL4* gene—mutations with functional consequences for either mode of repression should correspond to changes in NCOR binding if the effects are mediated by NCOR recruitment.

*ANGPTL4* expression was monitored by RT-qPCR in the absence (after treatment with the solvent) or the presence of a ligand. As ligands, we used either the inverse agonist PT-S264 or the agonist L165,041. Importantly, the ligands did not modulate *ANGPTL4* expression in PPARβ/δ knockout cells transduced with the empty vector or the DNA binding-deficient PPARβ/δ C91A-E92A mutant (supp. fig. S9 and fig. 5B). An extended description of the screen is available in the supplementary material including expression data for all mutants (supplementary appendices A–D) and a simplified overview with a classification of the effect of each mutant (supp. table S3).

**Figure 5.**
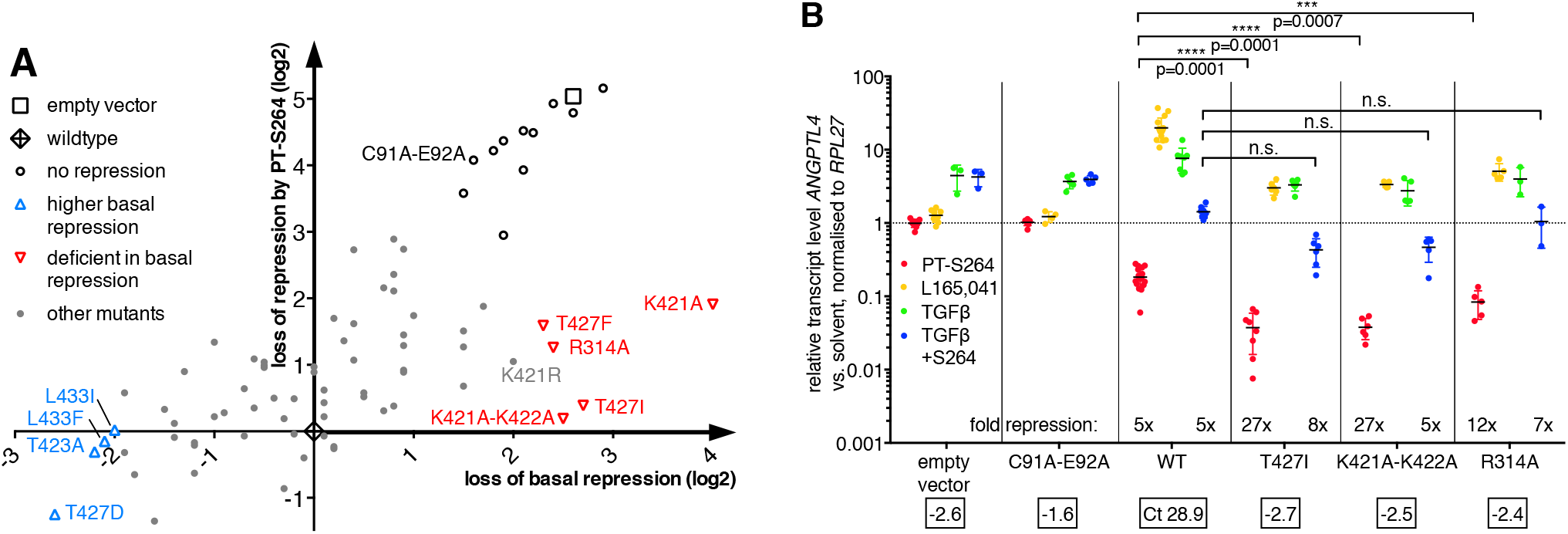
A retroviral reconstitution screen identifies PPARβ/δ mutants deficient in basal repression and response to PT-S264. MDA-MB231-*luc2* 2B3 *PPARD* KO cells stably expressing the murine ecotropic receptor were transduced with retroviruses derived from the empty vector or carrying *PPARD* cDNAs. The wildtype cDNA and a panel of 80 mutants (N between 1 and 8 per construct) were used. *ANGPTL4* expression was measured by RT-qPCR after treatment for 6 h. Values were normalised to the *RPL27* transcript. **A**: Effects on basal repression and PT-S264–dependent repression. *ANGPTL4* expression was plotted on the x axis as ΔΔCt(basal wildtype – basal mutant) vs. ΔΔCt(PT-S264 wildtype – PTS264 mutant) on the y axis. Open triangles denote mutants with modified basal repression (arbitrary cutoff of two PCR cycles relative to wildtype). Mean Ct values relative to the Ct value obtained from cells expressing the wildtype (WT) *PPARD* cDNA (N=20, mean Ct 28.9) were used. **B**: The relative expression of *ANGPTL4* was measured in cells transduced with the empty vector (N≥3), a retrovirus carrying the C91A-E92A DNA binding mutant *PPARD* cDNA (N≥4), the wildtype (N≥8), the T427I (N≥6), the K421A-K422A (N≥5), or the R314A mutant (N≥3) cDNAs and treated as indicated. The fold change relative to the solvent control sample was plotted. Significance levels were calculated *via* unpaired *t* tests between the fold repression values.

Briefly, our efforts identified several mutants which are generally compromised for receptor function, mutants which are compromised for ligand binding, mutants which show enhanced basal repression, and mutants which are deficient in basal repression. In this study, special interest pertains to mutants which affect repression. For visualisation of effects on repression, we plotted the ratio of basal *ANGPTL4* expression on the x axis as ΔΔCt(basal wildtype – basal mutant); the ΔΔCt term is due to the use of “ΔCt” values normalised to the housekeeping transcript. This way, loss of repression is reflected in a positive value. ΔΔCt(PT-S264 wildtype – PT-S264 mutant) is plotted on the y axis. The reconstituted wildtype receptor is at (0;0), and mutations which compromise both modes of repression cluster with the empty vector control in the upper right area of the graph (fig. 5A).

Mutants modified only in basal repression cluster on or below the x axis (fig. 5A), whereas mutants with differential capability for PT-S264–dependent repression should cluster on the y axis. We identified both kinds of mutants; however, deficiency in PT-S264–dependent repression was accompanied by loss of response to L165,041 in the few mutants we identified, indicating that the effect is not specific for the inverse agonist since ligand binding is compromised (see supp. fig. S9).

Strikingly, mutants with deficient basal repression (R314A, K421A-K422A, K421A, T427F, and T427I) show enhanced repression in the presence of PT-S264. These do not markedly deviate from the x axis (fig. 5), indicating that, while basal repression is compromised, PT-S264 is able to repress *ANGPTL4* expression to similar levels as achieved in the presence of the wildtype receptor (see also supp. fig. S9). This implies that the observed fold repression is higher by the amount lost from basal repression (see below).

Conversely, in cells expressing mutants with enhanced basal repression (negative on the x axis; I327A, T423A, T427D, and L433F), little or no PT-S264–stimulated repression was measured (supp. fig. S9). Therefore, the mutations affecting basal repression we identified here as well as the inverse agonist itself presumably modulate the affinity of PPARβ/δ towards the same repressors. We however cannot rule out that mutants with enhanced basal repression recruit additional other repressors. Moreover, due to low *ANGPTL4* expression levels in cells expressing these mutants, we are running into detection problems.

In the following experiments, we focus on three mutants with deficient basal repression, K421-K422A, T427I, and R314A. In panel B of fig. 5, detailed *ANGPTL4* RT-qPCR data for these three mutants are shown. Basal repression of *ANGPTL4* is relieved in cells expressing these mutants, resulting in a higher fold repression in the presence of PT-S264, while fold induction by the agonist L165,041 is lowered. In the presence of the activating stimulus TGFβ, repression by PT-S264 is functional in cells expressing these mutants, albeit to an extent similar to cells transduced with wildtype PPARβ/δ.

The main finding of the functional screen is that loss of basal repression is fully compensated for by the inverse agonist PT-S264, resulting in higher fold repression. *Vice versa*, enhanced basal repression limits repression by PT-S264. This leads to the hypothesis that the same corepressor(s) mediate both basal and inverse agonist-dependent repression. The K421A-K422A, T427I, and R314A mutants chosen for further analysis predominantly show PT-S264–mediated repression and thus allow to investigate whether basal repression and PT-S264–dependent repression indeed depend on the same corepressor(s). Due to data obtained by mass spectrometry (table 1), we assume that repression is mediated by NCOR or both NCOR and SMRT.

### PPARβ/δ mutants deficient in basal repression show diminished NCOR and SMRT recruitment to chromatin and increased RNA polymerase II recruitment in the basal state

We next asked if the mutants deficient in basal repression indeed show differential binding of NCOR and SMRT as predicted. Our previous genome-wide studies identified PPARβ/δ binding sites and target genes in different cellular model systems (1, 2, 4, 14) including the MDA-MB231-*luc2* cell line used here (4). The genes strongly regulated by PPARβ/δ ligands in cell lines are *ANGPTL4, PDK4, ABCA1*, and *PLIN2*. ChIP-qPCR analyses reveal that binding of both NCOR and SMRT to PPARβ/δ-responsive elements of the genes *ANGPTL4, PDK4*, and *PLIN2* is markedly reduced in cells that express the T427I, K421A-K422A, and R314A mutants (fig. 6). However, in the presence of PT-S264, NCOR and SMRT binding is restored by the T427I, K421A-K422A, and R314A mutants. Essentially, the same observation was made for HDAC3 (supp. fig. S10). As an additional control, the presence of mutated PPARβ/δ and RXR as well as NCOR recruitment by PT-S264 at the *ANGPTL4* PPAR binding site was confirmed (supp. fig. S11). Moreover, in the absence of ligand, binding of RNAPII to the *ANGPTL4, PDK4, PLIN2*, and *ABCA1* target gene TSSs is strongly reduced in *PPARD* KO cells reconstituted with wildtype PPARβ/δ (fig. 7A–D) but not in cells expressing the T427I and K421A-K422A mutants. Strikingly, PT-S264 treatment leads to reduced RNAPII binding in cells expressing these mutants. Notably, the mode of ligand binding is not altered in the mutants since an additional mutation that occludes the ligand binding pocket abrogates ligand function (see supp. fig. S12). RNAPII binding at the TSS of *TSC22D3*, which is not a PPAR target gene, is neither affected by expression of the PPARβ/δ-encoding cDNAs nor by PT-S264 (fig. 7F). This finding shows that the PPARβ/δ mutants deficient in basal NCOR, SMRT, and HDAC3 binding allow for increased RNAPII occupancy in the absence activating ligands. Conversely, these mutants recruit NCOR, SMRT, and HDAC3 in the presence of the inverse agonist PT-S264, and thus RNAPII binding is strongly reduced. These observations indicate that NCOR and SMRT mediate both basal and PT-S264–dependent repression by limiting RNAPII binding to PPARβ/δ target genes.

**Figure 6.**
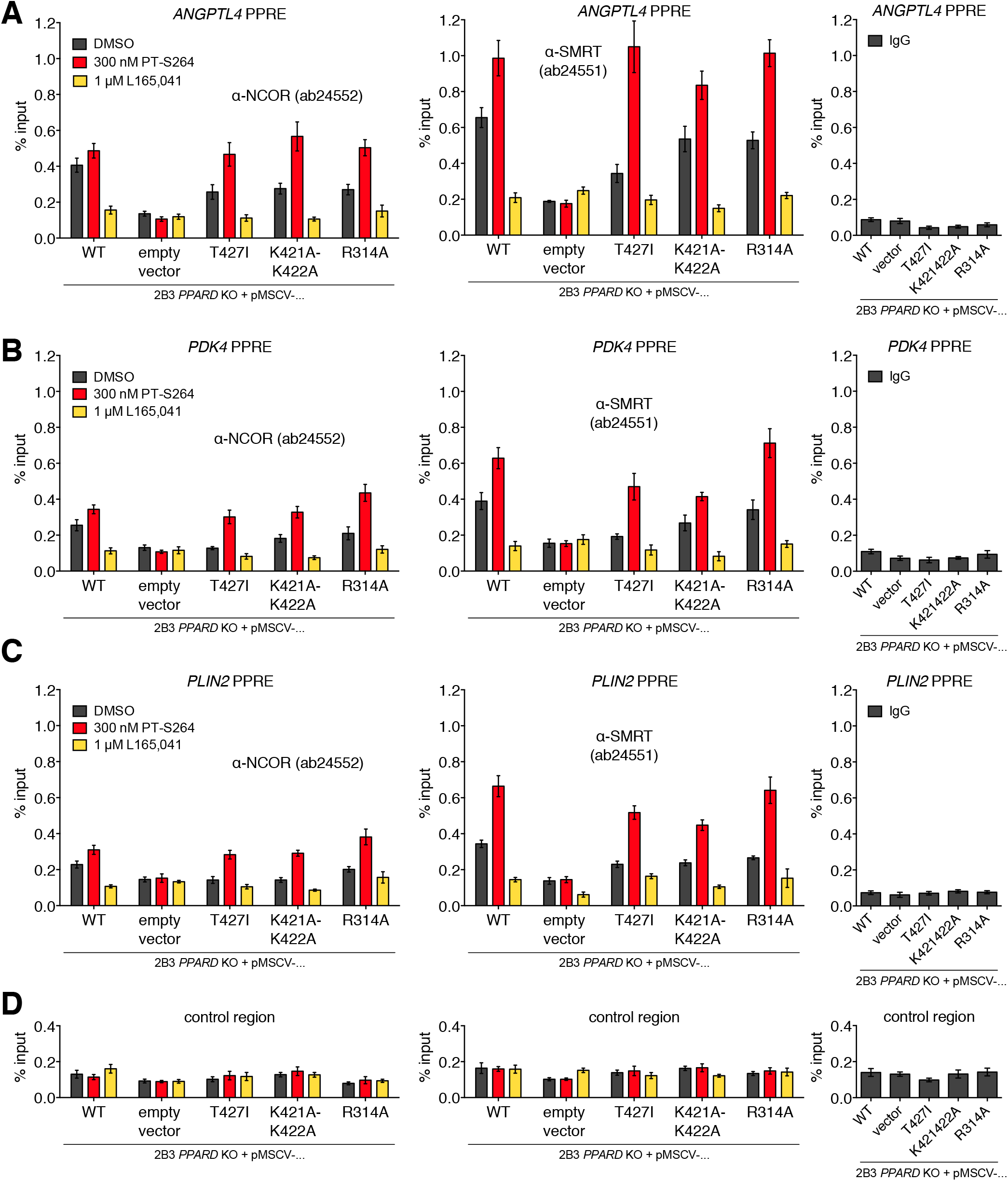
Corepressor recruitment by PPARβ/δ mutants deficient in basal repression. 2B3 *PPARD* KO cells transduced with the indicated constructs were treated with solvent, 1μM L165,041, or 300 nM PT-S264 for 30 min and subjected to ChIP-qPCR analysis using antibodies against NCOR or SMRT. The PPAR binding sites of *ANGPTL4* (+3,500 bp relative to the TSS, **A**), *PDK4* (–12,200 bp, **B**), *PLIN2* (–34,300 bp, **C**) and a negative control region (**D**) were amplified as indicated. Non-cognate IPs (IgG) were processed in parallel from the solvent-treated cells. Means and standard deviations of three technical replicates from a representative ChIP assay are plotted.

**Figure 7.**
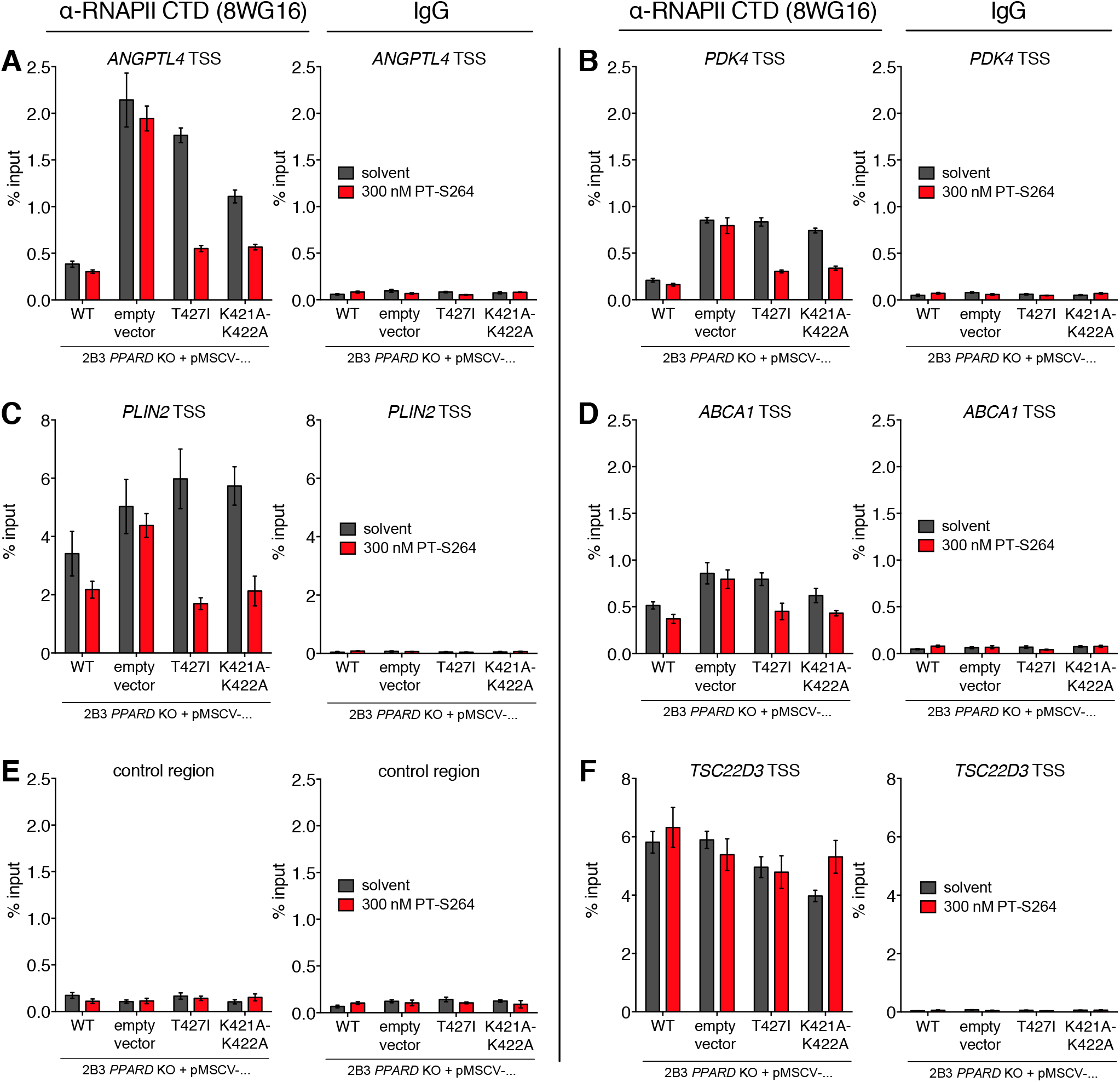
Regulation of RNA polymerase II binding by PPARβ/δ mutants deficient in basal repression. 2B3 *PPARD* KO cells transduced with the indicated constructs were treated with solvent or 300 nM PT-S264 for 30 min and subjected to ChIP-qPCR analysis using an antibody against RNA polymerase II. The TSS regions of *ANGPTL4* (**A**), *PDK4* (**B**), *PLIN2* (**C**), *ABCA1* (**D**), a negative control region (**E**), and the *TSC22D3* TSS (**F**) as an additional negative control were amplified as indicated. Means and standard deviations of three technical replicates from a representative ChIP assay are plotted.

### The role of deacetylase activity in repression by PT-S264

In order to investigate the role of the HDAC3 subunit of NCOR and SMRT complexes in repression of PPAR β/δ target genes, we treated cells with PT-S264 together with HDAC inhibitors. In wildtype MDA-MB231-*luc2*, neither TSA nor the HDAC3-selective compound apicidin abrogates repression of *ANGPTL4* elicited by PT-S264 (fig. 8A, upper left panel). However, repression of *ANGPTL4* is diminished by the HDAC inhibitors. PT-S264–mediated repression of *PDK4*, which is weaker in comparison to *ANGPTL4*, is also diminished by the HDAC inhibitors (fig. 8A, upper right panel), albeit to a lesser extent. Another PPARβ/δ target gene transcript, *PLIN2*, shows the same pattern (fig. 8A, lower left panel).

**Figure 8.**
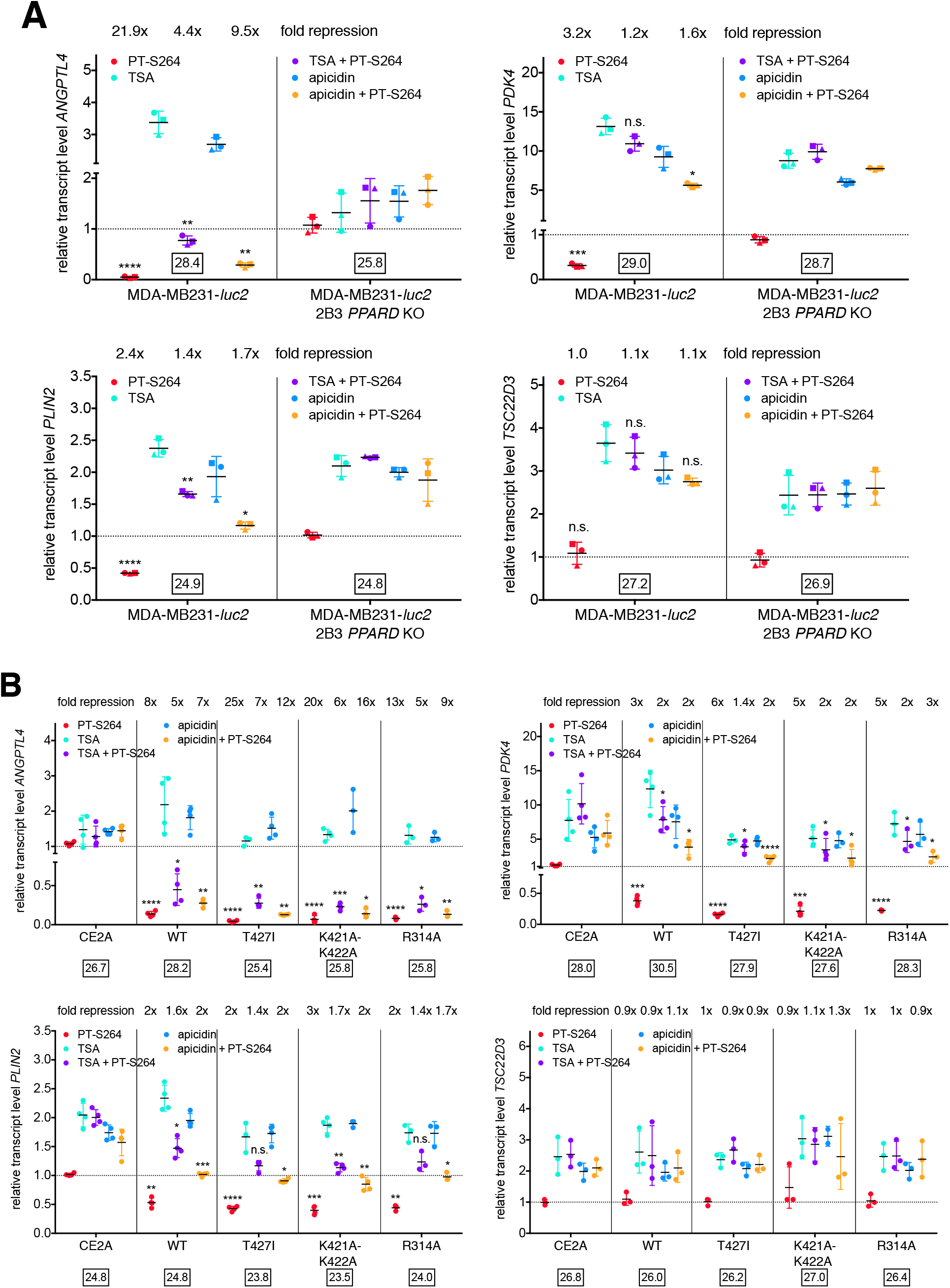
HDAC inhibitor sensitivity of PT-S264–mediated repression. Parental MDA-MB231-*luc2* and 2B3 *PPARD* KO cells (**A**) or 2B3 *PPARD* KO cells transduced with the indicated constructs (**B**) were treated with solvent, 300 nM PT-S264, 500 nM TSA, and 250 nM apicidin as indicated for 6 h, and RT-qPCR with primers against the *ANGPTL4, PDK4, PLIN2*, and *TSC22D3* transcripts was performed. Means and standard deviations of (A) n=3 independent experiments or (B) n=3–4 independent experiments are plotted. Mean normalised Ct values in the control condition are indicated. Significance levels were calculated *via* paired *t* tests. ****, p<0.0001; ***, p<0.001; **, p<0.01; *, p<0.05; n.s., not significant.

In KO cells with restored wildtype PPARβ/δ expression, PT-S264–mediated repression of *ANGPTL4* is insensitive to the HDAC inhibitors, whereas in *PPARD* KO cells expressing mutants deficient in basal repression, repression of *ANGPTL4* by PT-S264 is partially sensitive to HDAC inhibition (fig. 8B). The *PDK4* transcript is affected similarly. This clearly suggests that the NCOR and SMRT complexes recruited to PPARβ/δ target genes exert both deacetylase-dependent and deacetylase-independent repressive functions; the catalytic activity of HDAC3 contributes to but is not sufficient for full repression. This is in agreement with the observation that *ANGPTL4* expression is only weakly induced by HDAC inhibitors (fig. 8B and (4)).

In general, weaker repression in 2B3 cells reconstituted with the wildtype *PPARD* cDNA relative to the parental cells was observed (supp. fig. S6). We attribute this to sequestration of corepressors by overexpressed wildtype PPARβ/δ, which recruits NCOR and SMRT considerably stronger than the mutants deficient in basal repression (fig. 6). Sequestration might differentially affect the availability of free NCOR and SMRT.

## DISCUSSION

NCOR and SMRT figure in gene regulation by a multitude of transcription factors such as the glucocorticoid receptor (56, 57), other nuclear receptors (8, 58), POZ-containing, basic helix-loop-helix, and basic leucine zipper factors (59), the Notch effector RBP-J (60), and MECP2 (methylated CpG binding protein 2) (61). Therefore, elucidation of regulatory mechanisms used by NCOR and SMRT complexes is of great interest. We show here that PPARβ/δ inverse agonists interfere with recruitment of the transcription-permissive Mediator subunits MED1, MED13L, and MED26 as well as TFIIB and RNAPII. PPARβ/δ mutants deficient in NCOR and SMRT recruitment in the basal state allow for enhanced RNAPII binding in the absence of ligands, while the inverse agonist PT-S264 restores both NCOR and SMRT binding to these mutants and loss of RNAPII at target gene promoters. Repression is only partially sensitive to HDAC inhibition, which indicates that NCOR/SMRT complexes exert deacetylase-dependent and deacetylase-independent functions to restrain transcription of PPARβ/δ target genes.

HDAC-independent repression mechanisms exerted by NCOR/SMRT have also been reported in other contexts: (I) In mice, Ncor and Smrt complexes carry out essential functions that are independent of Hdac3 activity (62), and (II) HDAC-independent repression of human papillomavirus transcription and replication by NCOR was demonstrated (63). (III) Finally, whereas MECP2 represses transcription *via* an HDAC-independent mechanism (64), interaction with NCOR/SMRT is crucial for MECP2 function (61), and it was reported recently that the detrimental effects of Mecp2 overexpression in mice do not depend on the catalytic activity of Hdac3 (65).

Our data suggest that deacetylation by HDAC3 contributes to but is not sufficient for basal repression of PPARβ/δ target genes. Thus, an additional enzymatic function of HDAC3-containing complexes or, alternatively, inhibition of an activator-borne enzymatic activity may be required for efficient silencing of gene expression. We speculate that acetyltransferases are inhibited by NCOR and SMRT complexes since (i) acetyltransferase activity is required for transcription in nucleosome-free systems (66), (ii) GTFs that contribute to reinitiation—TFIIB, TFIIE, and TFIIF—are acetylated *in vivo* (67, 68), and (iii) the acetyltransferase activities of CBP and p300 are inhibited by recombinant purified fragments of NCOR and SMRT *in vitro* (69, 70). Moreover, Mediator interacts with the acetyltransferase-bearing SAGA complex, and both Mediator and SAGA are necessary for RNAPII-mediated transcription in yeast (71, 72, 73). Partial sensitivity of PT-S264–dependent repression to HDAC inhibitors (fig. 8) strongly suggests that NCOR and SMRT recruited by PPARβ/δ use an additional mechanism to restrict activator function. It is striking that, in cells ectopically expressing PPARβ/δ mutants with deficient basal repression, PT-S264–dependent repression is more sensitive to HDAC inhibition than in cells ectopically expressing the wildtype receptor (fig. 8B). This finding suggests that the NCOR/SMRT complexes involved in basal repression, whose binding is restored by PT-S264 in cells expressing the mutants (fig. 6), rely on HDAC activity to a larger extent compared to the complexes that are recruited to the wildtype receptor upon treatment with PT-S264.

Induction of the PPARβ/δ target gene *ANGPTL4* by TGFβ depends on SMAD3 (18), which interacts with CBP and p300 to activate transcription (74). CBP and p300 are also paramount coactivators of NFκB (75) and AP-1 (76) transcription factors, which are major targets of transrepression by nuclear receptors (57, 76, 77, 78). These data underscore a possible role for acetyltransferase inhibition in gene regulation by NCOR and SMRT. Since deacetylation and inhibition of acetyltransferase activity act towards the same outcome, the two functions may conceivably act in parallel for more efficient suppression of transcription. Another possibility would be that ubiquitin ligase and deubiquitinase activities which reside in subunits of HDAC3-containing complexes contribute to deacetylase-independent repression. Ubiquitination by TBLR1 (8) could commit activators for degradation, or H2B deubiquitination by USP44 might figure in repression of *ANGPTL4* by NCOR/SMRT (79).

Mediator supports TFIIB binding to promoters, and both Mediator and TFIIB facilitate recruitment of RNAPII to promoter-bound GTFs (30). This notion is in agreement with both older and more recent studies in yeast (43, 71, 80, 81), murine (82), and human cells (83) which indicate concurrent interactions of Mediator, TFIIB, and RNAPII. MS data from *S.cerevisiae* and from human cells identify RNAPII, TFIIB, and TFIIF among the strongest interactors of Mediator (71, 83), and these factors are necessary for reinitiation (25). Notably, another recent study revealed that human genes which require *de novo* RNAPII recruitment for induction of transcription depend on TFIIB availability (84). It is unclear how transcription factors modulate reinitiation, and *in vivo* evidence of reinitiation and scaffold complexes is lacking (32). Our ChIP data (fig. 1 and supp. fig. S2), which demonstrate impairment of TFIIB and RNAPII binding by PPARβ/δ inverse agonists, are in agreement with findings obtained from *in vitro* systems that describe recruitment of RNAPII and TFIIB (25, 30) to scaffold factors and thus may represent *in vivo* correlates of reinitiation and scaffold complexes. RNAPII binding to the *ANGPTL4* promoter is affected similarly by PT-S264 and the TFIIH-XPB inhibitor (85) triptolide (fig. 1A); triptolide is expected to affect both initiation and reinitiation. Our ChIP data do not allow for discrimination between the first and subsequent rounds of transcription; hence, we cannot conclude whether the first round of transcription is affected as well.

Our observations furthermore suggest that TFIIB and RNAPII recruitment coincides with the presence of MED1, MED13L, and MED26 (fig. 2). This notion is supported by previous work of others which identified MED1 (44, 45), MED13L (48), and MED26 (48, 86, 87) as subunits that are predominantly present in RNAPII-associated Mediator. Our observations are compatible with the model that the MED13-containing kinase module blocks RNAPII association with Mediator (31) since MED26 and MED13 binding is mutually exclusive (48). Notably, co-occurrence of MED13L and MED26 was described (44, 48, 86, 87). Our model thus favours an RNAPII-associated Mediator state which harbours the MED1, MED13L, and MED26 subunits, and this is counteracted by PT-S264 at PPARβ/δ target genes.

It should be noted that we cannot formally exclude involvement of corepressors other than NCOR and SMRT in PT-S264–dependent repression. Genetic deletion of *NCOR1* was unsuccessful in our hands; this is consistent with the notions that NCOR function may be necessary for regulation of the cell cycle (88), genome stability (89), or other critical processes (58). Cellular models without this limitation or knock-in approaches might be suitable to resolve whether NCOR and SMRT complexes are sufficient for repression by PPARβ/δ inverse agonists. Modification of endogenous coding sequences would also circumvent possible effects of overexpression such as cofactor sequestration. Notably, after reconstitution of PPARβ/δ expression, PT-S264–dependent repression is largely insensitive to HDAC inhibition, which is in contrast to partial sensitivity observed in the parental cell line. Moreover, HDAC inhibition leads to weaker upregulation of PPARβ/δ target genes in the parental cells compared to cells with reconstituted PPARβ/δ. This could be due to higher recruitment of SMRT relative to recruitment of NCOR after reconstitution of PPARβ/δ expression (fig. 6), whereas the ratio of ChIP-qPCR signals obtained with the same antibodies is closer to one in wildtype cells (fig. 3). However, the preferential dependence of SMRT complexes on deacetylase activity relative to NCOR complexes is highly speculative.

Taken together, we propose that PPARβ/δ recruits NCOR and SMRT, either of which or both block MED1, MED13L, and MED26 recruitment to promoter-bound GTFs *via* both deacetylase-dependent and -independent functions to interfere with TFIIB and RNAPII binding. A model depicted in fig. 9 summarises our conclusions. Additional insight will require identification of the subunits and domains of NCOR and SMRT complexes which exert deacetylase-independent repression and their target proteins, and it will be interesting to probe possible differential functions of NCOR and SMRT.

**Figure 9.**
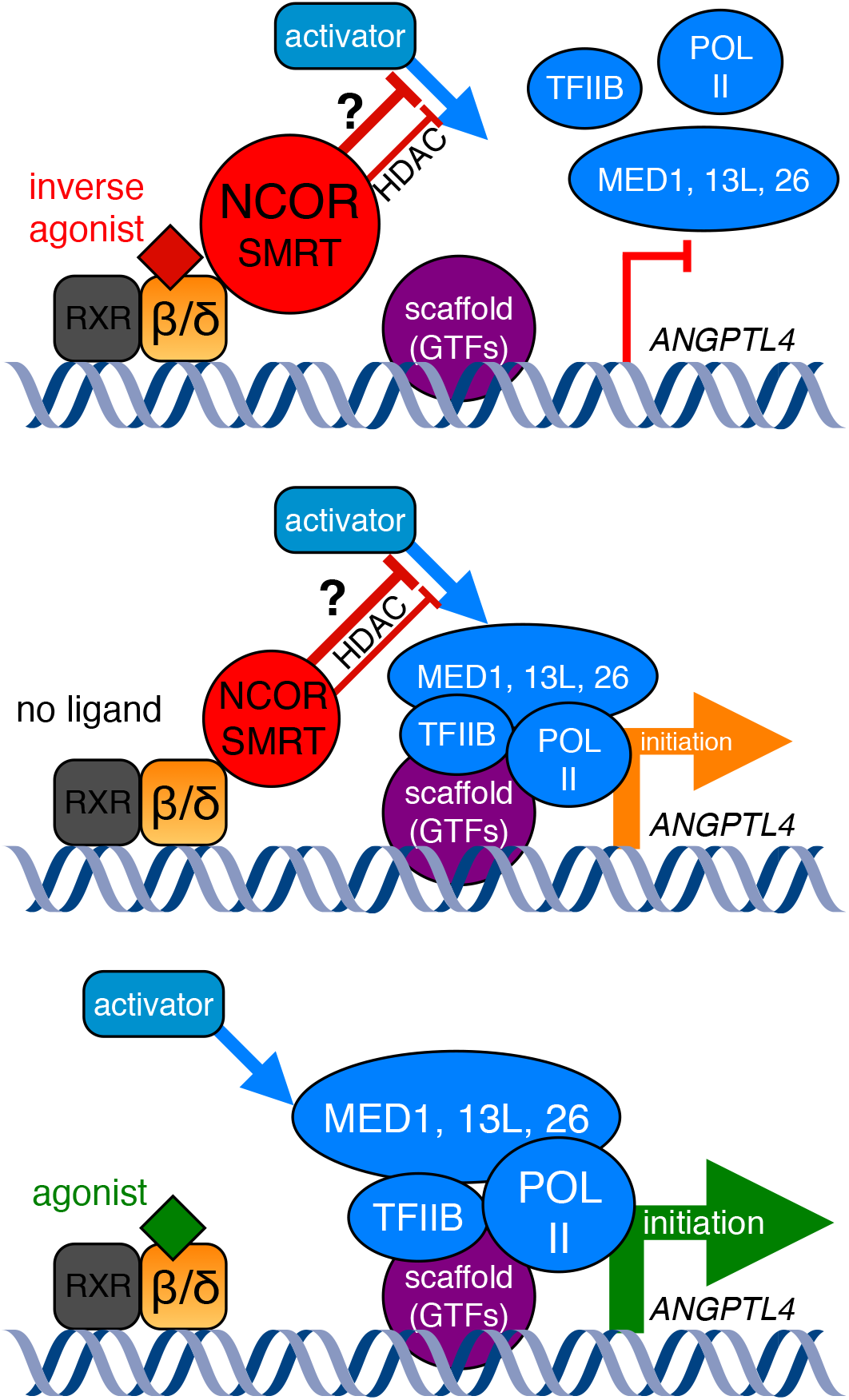
Ligand-dependent regulation of *ANGPTL4* transcription initiation by PPARβ/δ and NCOR/SMRT. Data from this study show that the PPARβ/δ inverse agonist PT-S264 interferes with activator-stimulated recruitment of MED1, MED13L, MED26, TFIIB, and RNAPII. Mutations of PPARβ/δ which abrogate NCOR and SMRT binding in the basal state allow for increased binding of RNAPII, and NCOR and SMRT binding is restored upon addition of PT-S264. The latter observations were made at other PPAR target genes as well. Taken together with our previous observation that synthetic PPAR agonists together with other activating stimuli synergistically induce *ANGPTL4* transcription (18), we propose the model that basal and ligand-dependent repressor recruitment limit the ability of activators to induce transcription reinitiation (and possibly initiation) *via* NCOR, SMRT, or both. The corepressors use both deacetylase-dependent and deacetylase-independent mechanisms, the latter of which are not elucidated as of now.

## DATA AVAILABILITY

The mass spectrometry proteomics data have been deposited to the ProteomeXchange Consortium via the PRIDE (90) partner repository with the dataset identifier PXD012818 and 10.6019/PXD012818.

## SUPPLEMENTARY DATA

Twelve supplementary figures, three supplementary tables, an extended description and interpretation of the mutant screen, and an appendix with expression values for all mutants is available for download as a PDF file.

## ACKNOWLEDGEMENTS

We gratefully acknowledge the performance of RIME assays by Garrett Shafer (Active Motif, Inc.) and the help of Matthias Spiller-Becker (Active Motif, Inc.), Tiffany Yen (Active Motif, Inc.), and Florian Finkernagel with mass spectrometry data. We thank Guntram Suske, Ho-Ryun Chung, Nathalie Hoffmann, and Florian Finkernagel for helpful discussions and critical reading of the manuscript.

## AUTHOR CONTRIBUTIONS

Conceived, designed, and performed the experiments: NL TA. NL performed the vast majority of the experiments. Analysed data: NL BF TA. Generated and contributed reagents and materials: NL CLB SZ BW MD WED TA. Supervised the study and wrote the paper: TA.

