## Supplementary information file for "PPARβ/δ recruits NCOR and regulates transcription reinitiation"

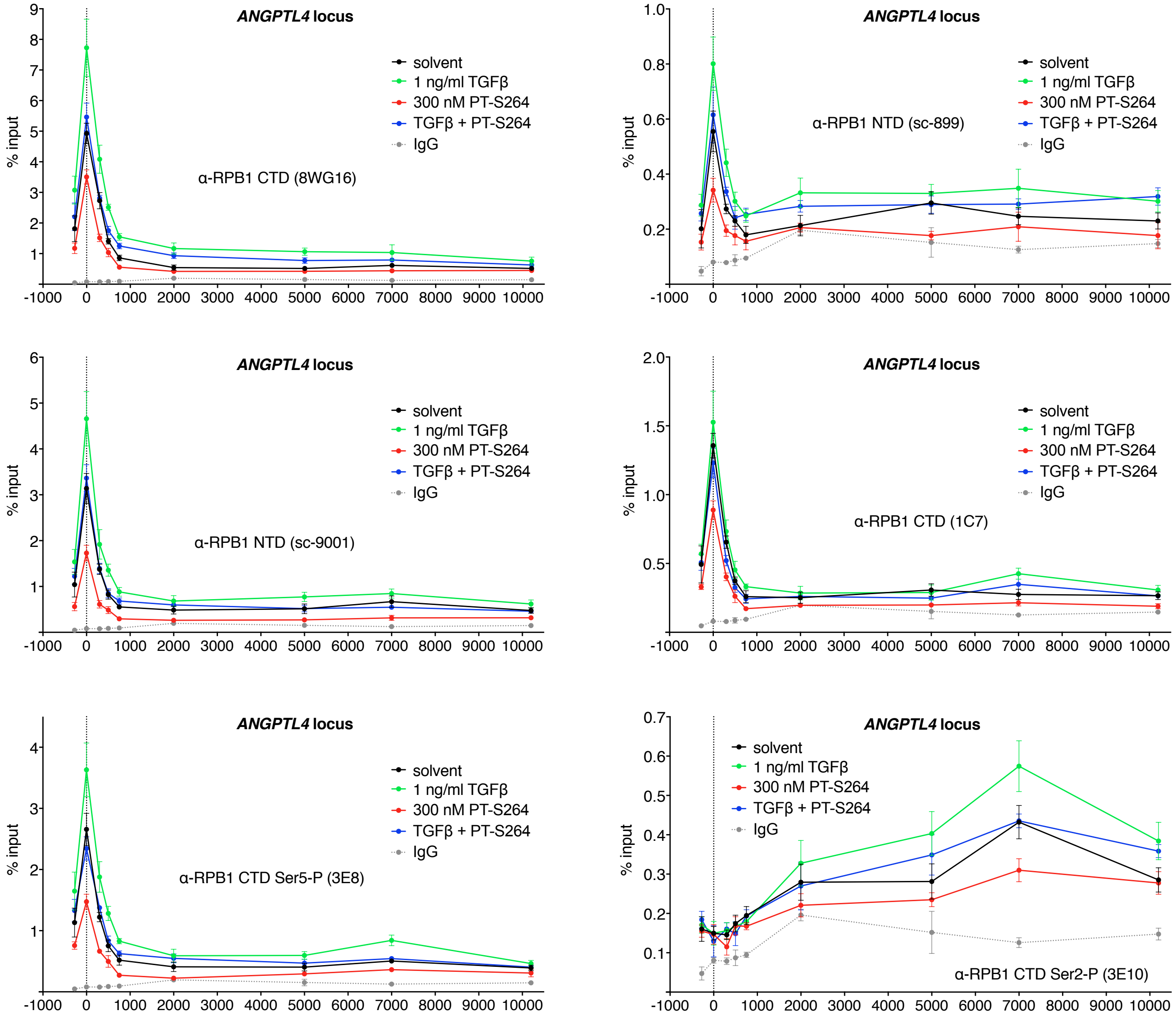


**Supp. fig. S1. RNA polymerase II scanning ChIP at the *ANGPTL4* locus in Caki-1 cells.** Caki-1 cells were treated for 30 min as indicated and subjected to ChIP-qPCR analysis with antibodies against RNA polymerase II large subunit (RPB1) N-terminal or C-terminal domain (NTD/CTD), or phosphorylated forms of the CTD, as indicated. Primer pairs across the promoter and the entire coding region were used. A sample obtained with an irrelevant IgG pool from non-immunised rabbits was processed in parallel. Terms in parentheses denote the ordering numbers of the antibodies.

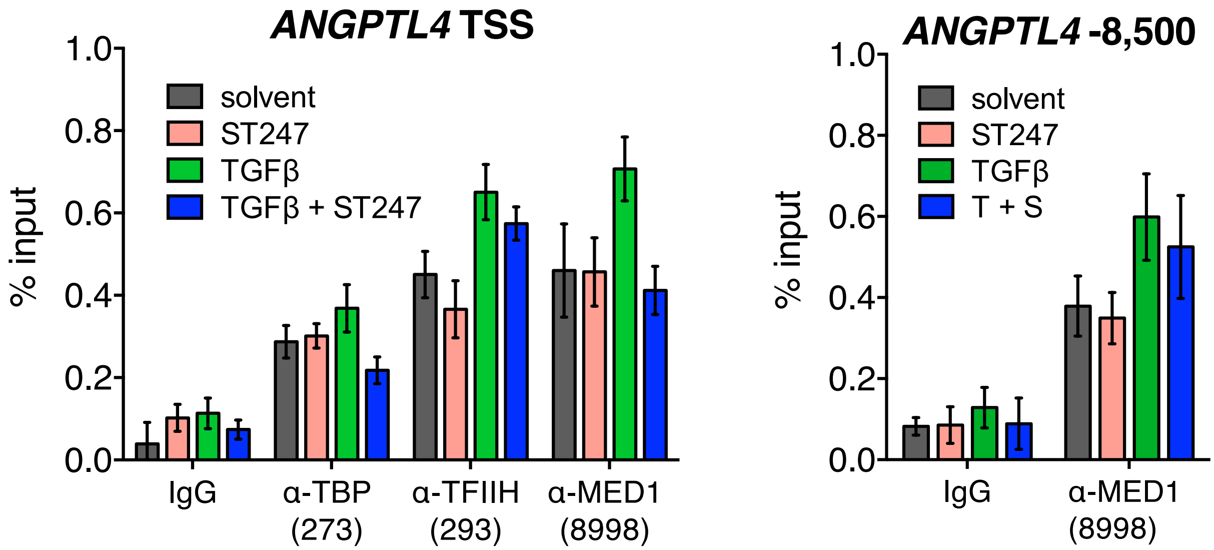


**Supp. fig. S2. Binding of TBP, TFIIH, and MED1 at the *ANGPTL4* TSS and the TGFβ- responsive enhancer** (8,500 bp upstream of the TSS) in the presence of TGFβ, ST247, or both. Caki-1 cells were treated as indicated (T=TGFβ, 2 ng/ml; S=ST247, 300 nM) for 30 min. ChIP-qPCR analyses were performed with primer pairs amplifying the *ANGPTL4* TSS (left panel) or TGFβ-responsive enhancer (right panel, 8,500 bp upstream of the TSS).

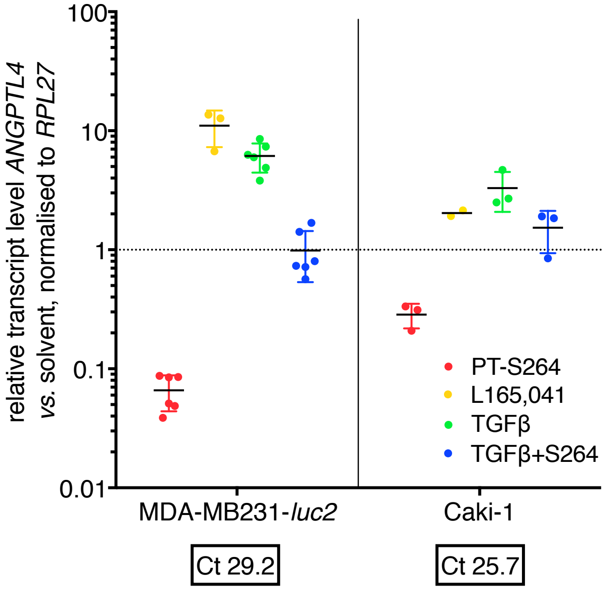


**Supp. fig. S3. Regulation of *ANGPTL4* by PPARβ/δ ligands in MDA-MB231-*luc2* and Caki-1 cells.** After treatment of the cells for 6 h with the indicated stimuli (PT-S264, 300 nM; L165,041, 1 µM; TGFβ, 2 ng/ml), RT-qPCR analysis of *ANGPTL4* expression was performed using the *RPL27* transcript for normalisation. N≥2 as indicated; the error bars represent standard deviations calculated from the analyses of biological replicates. Mean basal Ct values are indicated.

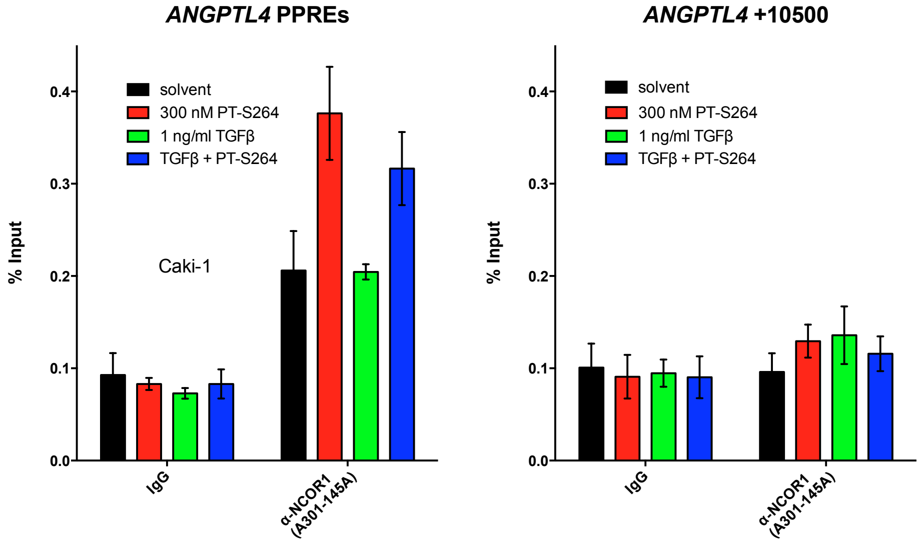


**Supp. fig. S4. NCOR recruitment to the *ANGPTL4* PPAR binding site in Caki-1 cells.** Near-confluent cultures were treated for 30 min as indicated, and ChIP-qPCR analyses were performed with an NCOR-specific antibody and an irrelevant IgG pool from non-immunised rabbits.

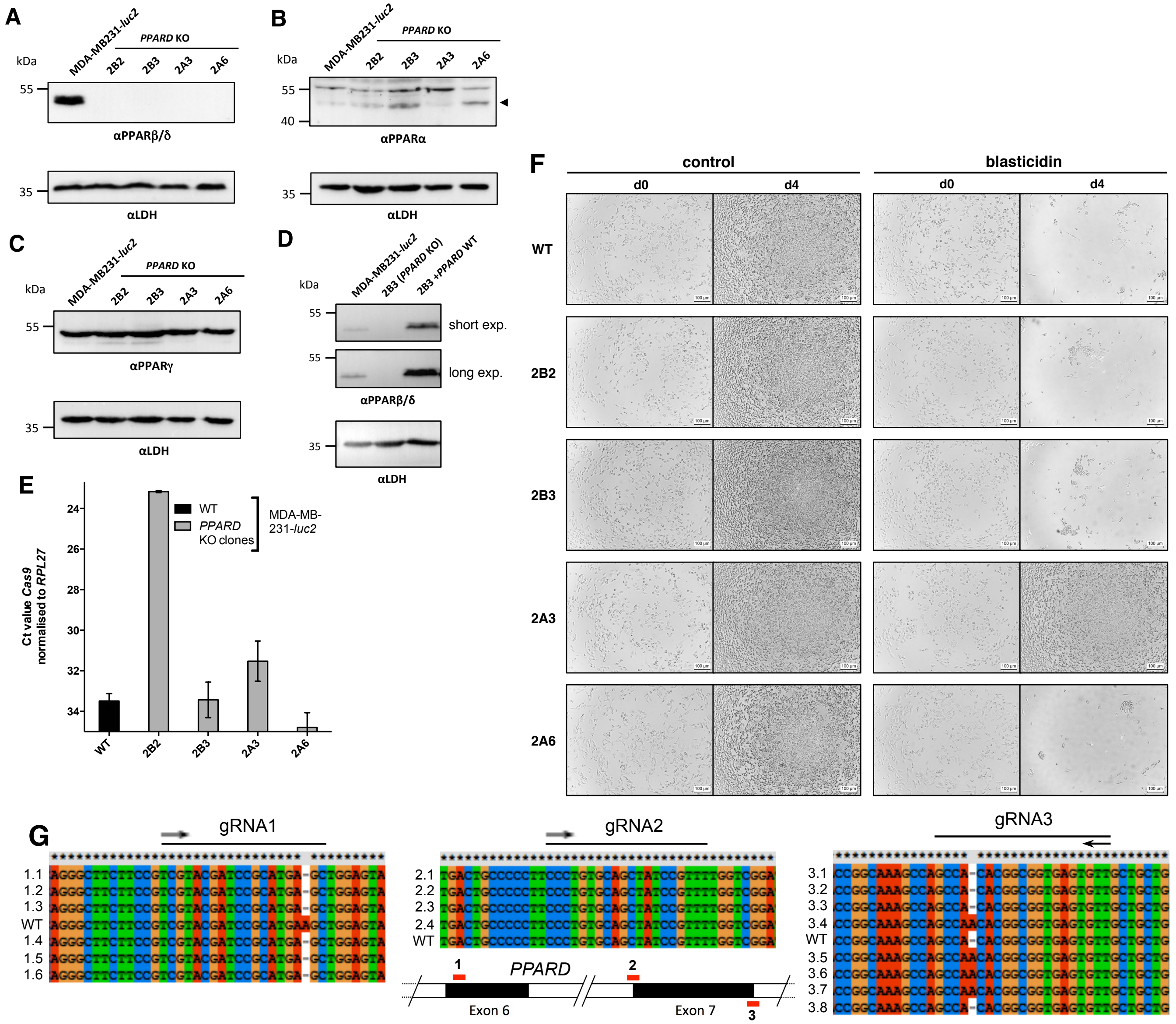

**Supp. fig. S5.** Characterisation of *PPARD* knockout clones and reconstitution of PPARβ/δ expression. **A**: Immunoblot analysis of PPARβ/δ expression in the parental MDA-MB231-*luc2* cell line and four knockout clones (2B2, 2B3, 2A3, 2A6). The samples were probed for LDH (lactate dehydrogenase) expression as a loading control. **B,C**: Analysis of PPARα (**B**) and PPARγ (**C**) expression in the KO clones. The arrowhead in **B** indicates the band at the expected molecular weight for PPARα. **D**: Reconstitution of the 2B3 *PPARD* KO clone. 2B3 cells were stably transfected with an expression vector for the murine ecotropic receptor and subsequently transduced with a *PPARD*-encoding retrovirus. Two different exposure times are shown for the PPARβ/δ immunoblot. **E**: RT-qPCR analysis of *Cas9* expression in the KO clones. **F**: Blasticidin resistance test of the KO clones. Clone 2A3 stably integrated the resistance marker. **G**: Sequencing data of vectors carrying cloned PCR fragments derived from genomic DNA of clone 2B3 (TOPO-TA–based cloning). Primer pairs amplifying genomic fragments around the target sites of guide RNAs 1, 2, and 3 were used, and the obtained sequencing results were aligned to the wildtype genomic sequence using ClustalX. For reference, a map of exons 6 and 7 of the *PPARD* locus and the expected binding sites of the gRNAs is shown.

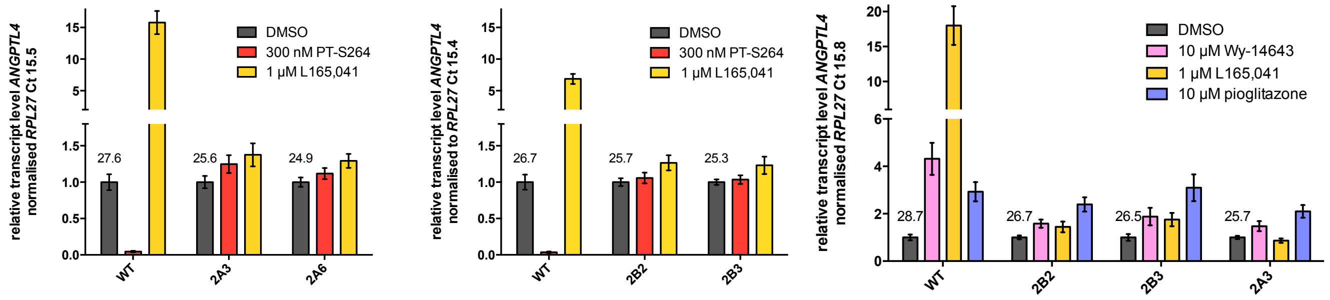


**Supp. fig. S6. Regulation of *ANGPTL4* transcription by PPAR ligands in PPARβ/δ knockout clones.** The parental wildtype (WT) MDA-MB231-*luc2* cells or the indicated PPARβ/δ knockout clones were subjected to RT-qPCR analyses of *ANGPTL4* expression using the *RPL27* transcript for normalisation. The cells were treated for 6 h as indicated. A representative experiment is shown. Error bars represent standard deviations calculated from three technical replicates. Mean basal Ct values are included in the graphs.

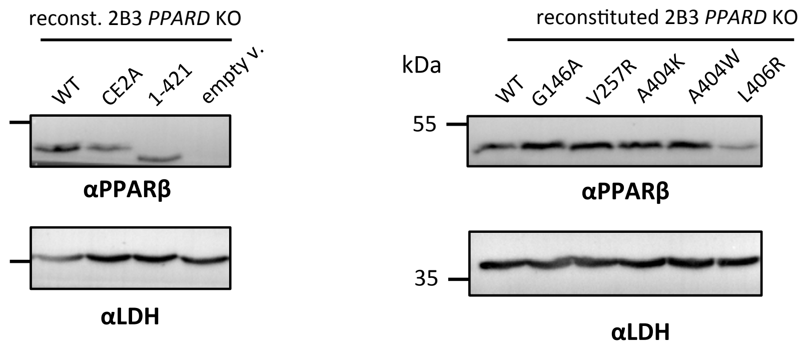


**Supp. fig. S7. Expression of PPARβ/δ mutants in the 2B3 knockout clone.** 2B3 cells stably expressing the murine ecotropic receptor were transduced with retroviruses derived from the empty vector or carrying *PPARD* cDNAs. PPARβ/δ expression was measured by immunoblotting. CE2A, abbreviation for C91A-E92A. 1–421: ΔL-box, ΔAF2 (residues 422–441 are deleted). The LDH immunoblot serves as a loading control.

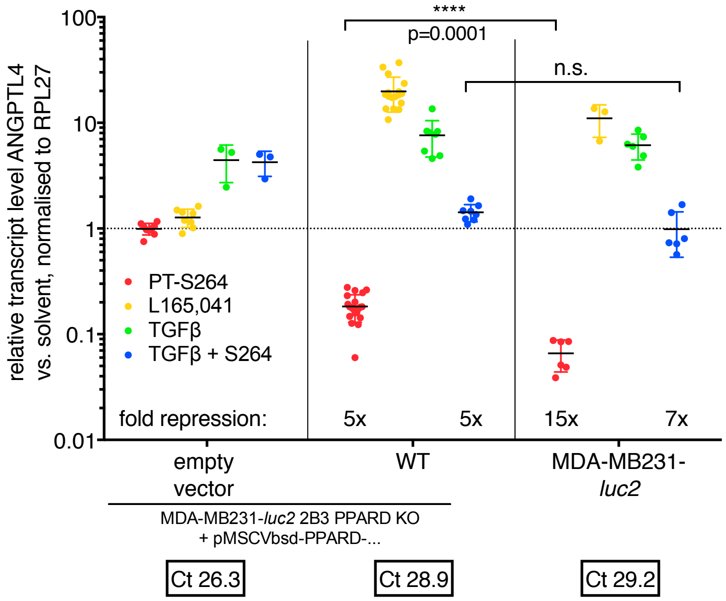


**Supp. fig. S8. Repression of *ANGPTL4* transcription by PPARβ/δ ligands in the PPARβ/δ knockout clone 2B3 transduced with the empty vector (left panel), in the 2B3 PPARβ/δ knockout clone transduced with the wildtype *PPARD* cDNA (WT), and in the parental MDA-MB231-*luc2* cells (right panel, same as in fig. S3).** After treatment of the cells for 6 h with the indicated stimuli (PT-S264, 300 nM; L165,041, 1 µM; TGFβ, 2 ng/ml), RT-qPCR analysis of *ANGPTL4* expression was performed using the *RPL27* transcript for normalisation. N≥3 as indicated; the error bars represent standard deviations calculated from the analyses of biological replicates. Significance values were calculated using unpaired *t* tests. ****, $p < 0.0001$; n.s., not significant. Mean basal Ct values are indicated.

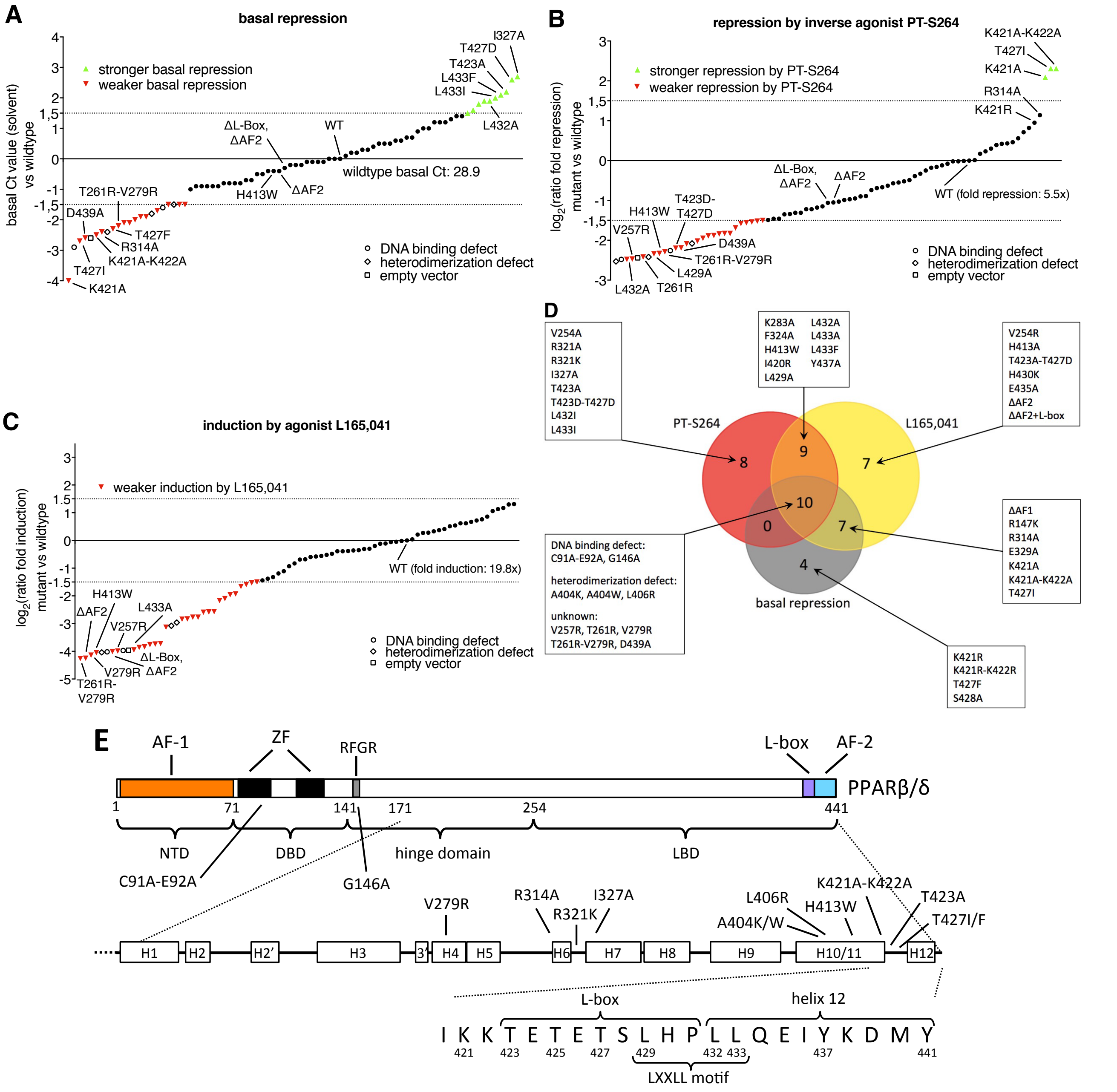


**Supp. fig. S9.** A retroviral reconstitution screen identifies PPARβ/δ mutants deficient in ligand response and basal repression. 2B3 cells stably expressing the murine ecotropic receptor were transduced with retroviruses derived from the empty vector or carrying *PPARD* cDNAs. The wildtype cDNA and a panel of 80 mutants (N between 1 and 8 per construct) were used. *ANGPTL4* expression was measured by RT-qPCR after treatment for 6 h. The *RPL27* transcript was used for normalisation. **A**: Effects on basal repression. Mean Ct values relative to the Ct value obtained from cells expressing the wildtype (WT) *PPARD* cDNA (n=20, mean Ct 28.9) were plotted. **B,C**: Effects on repression elicited by 300 nM PT-S264 and activation of transcription by 1 µM L165,041. The $\log_2$ ratio (ΔΔΔCt) of the relative expression in the respective mutant to the WT was plotted. The mean relative expression (fold change) to the solvent control in cells expressing the WT cDNA is indicated. **D**: A Venn diagram representing overlapping deficiencies of the indicated mutants in PT-S264–dependent repression, basal repression, and L165,041–dependent induction of *ANGPTL4* expression. Mutants deficient in all three PPARβ/δ-dependent effects (lower left box) are presumably generally defective. **E**: A schematic representation of the PPARβ/δ protein including structural features, sequence motifs, and the locations of selected mutations.

### Detailed results and discussion of the PPARβ/δ mutants screening

Inherent technical and biological variation of the RT-qPCR–based readouts hampered direct comparison of cohorts of cells infected at different time points. Therefore, we calculated the *ANGPTL4* mRNA expression levels relative to the mean of twenty biological replicates from wildtype PPARβ/δ-reconstituted 2B3 cells. We plotted the $\log_2$ fold change mutant / fold change wildtype and used a cutoff of $2^{1.5}$, equivalent to 2.8 fold (supp. fig. S9). The positions of selected mutations referred to below are highlighted in fig. S9E. Our screen identified six types of mutations:

(i) Mutations which disrupt basal repression while the receptor is still otherwise functional (fig. S9A). These are K421A-K422A, K421A, and T427I; in cells expressing these mutants, ligand function is not generally impaired (supp. figs. S9B–C, see below; this established that these receptor mutants bind to DNA and allow for ligand binding), while *ANGPTL4* expression is not downregulated in the absence of either ligand. The R314A, T427F, and K421R mutants are also deficient in basal repression but scored below the arbitrary cutoff chosen here when analysing PT-S264–mediated repression (supp. fig. S9B, see below).

(ii) Mutations which enhance basal repression (supp. fig. S9A). The function of the ligands L165,041 and PT-S264 is not generally impaired; the I327A, T423A, T427D, L432A, L433F, and L433I mutants cause diminished *ANGPTL4* expression in the absence of ligand compared to wildtype PPARβ/δ.

(iii) Function of the inverse agonist PT-S264 is deficient upon expression of the R321K, T423D-T427D, L429A, and L433A mutants, while basal repression is mostly unaffected (supp. fig. S9B).

(iv) Function of the agonist L165,041 is severely impaired upon expression of LxxLL, AF-2, Y437A, and C-terminal deletion mutants (ΔAF-2 and ΔL-box–AF-2); see supp. fig. S9C.

(v) Ligand binding is presumably affected by the F324A and H413W mutations (supp. fig. S9B,C). These amino acid residues face inside the ligand binding pocket (inferred from structural data, PDB ID 3TKM).

(vi) Mutations which lead to general defects of PPARβ/δ function (supp. fig. S9A–C); these affect DNA binding (C91A-E92A and G146A), RXR heterodimerisation (A404K, A404W, and L406R; inferred from structural data, PDB ID 3DZY), and coregulator interaction in general (V257R, T261R, V279R). The D439A mutation might preclude engagement of the agonist-induced conformation of H12. However, we cannot exclude that D439A, T261R, and V279R negatively influence protein stability, as it was observed for L406R (supp. fig. S7) or that the V257R, T261R, V279R, or D439A mutations unexpectedly impair chromatin binding of the mutated receptor.

Taken together, the screening system showed the predicted results in all control mutants, identified mutants deficient in basal and PT-S264–dependent repression, and yielded some unexpected findings: Deletion of helix 12 (AF2) and its connecting loop (L-box) did not abrogate PT-S264–mediated downregulation of *ANGPTL4* expression (supp. fig. S9B), demonstrating that the AF2 domain is not required for repression by the inverse agonist. Loss of basal repression in cells expressing the K421A, K421A-K422A, and T427I mutants leads to augmented repression in the presence of the inverse agonist (supp. fig. S9B), resulting from a higher basal transcription. Another striking observation pertains to the role of the synthetic agonist L165,041: The mutants with deficient basal repression show only weak upregulation of *ANGPTL4* after agonist exposure (supp. fig. S9B). In the presence of TGFβ, PT-S264–dependent repression in cells expressing these mutants is slightly enhanced (fig. 5D). Induction by the agonist is again weaker upon expression of the mutants. This indicates that the main function of L165,041 is to relieve basal repression in this cellular model system. Taken together, we conclude that the agonist relieves basal repression by decreasing affinity towards corepressors, and the inverse agonist presumably increases affinity towards corepressors and hence leads to stronger repression. In the agonist, bound conformation, helix 12 can apparently block the binding surface used by corepressors, explaining why it is expendable for their recruitment.

Supplementary table 1 on the following page provides a classification of each mutant on *ANGPTL4* expression regarding ligand functions and basal repression. Appendices A–D contain relative expression values for all mutants and $\log_2$ fold changes relative to mean values of samples from cells expressing the wildtype cDNA.

To test whether PT-S264 binds inside the ligand binding pocket of the mutants, as we showed previously for the wildtype LBD using a commercially available displacement assay (Toth *et al.* 2016), we made use of the finding that the H413W mutation presumably obstructs binding of any ligand (see supp. fig. S9). H413W puts a bulky residue inside the pocket of the LBD and, as a single mutation, has no overt effect on basal repression, which is preserved. Therefore, we introduced the H413W mutation in addition to the T427I, K421A-K422A, and R314A mutations to clarify the question whether the inverse agonist binds to the ligand binding pocket of the mutants. RT-qPCR data obtained with the dual mutations H413W-K421A-K422A, H413W-T427I, and H413W-R314A (see supp. fig. S12) show that the H413W mutation restores basal repression when introduced together with either K421A-K421A or T427I, which strongly suggests that H413W here enforces a conformation of the protein which overrides the negative effect of the other mutations on corepressor binding in the basal state. In contrast, the effect on basal repression of the R314A mutation, which unlike the other mutant residues is not localised in helix 11 of the LBD, was unaffected (supp. fig. S12); formally, it is possible that the H413W-R314A mutant is not expressed or does not bind to DNA). As predicted, all three mutants carrying the additional H413W mutation are not responsive to both L165,041 and PT-S264. This indicates that ligand binding is compromised by the H413W mutation and therefore points to binding of PT-S264 inside the LBDs of the mutants.

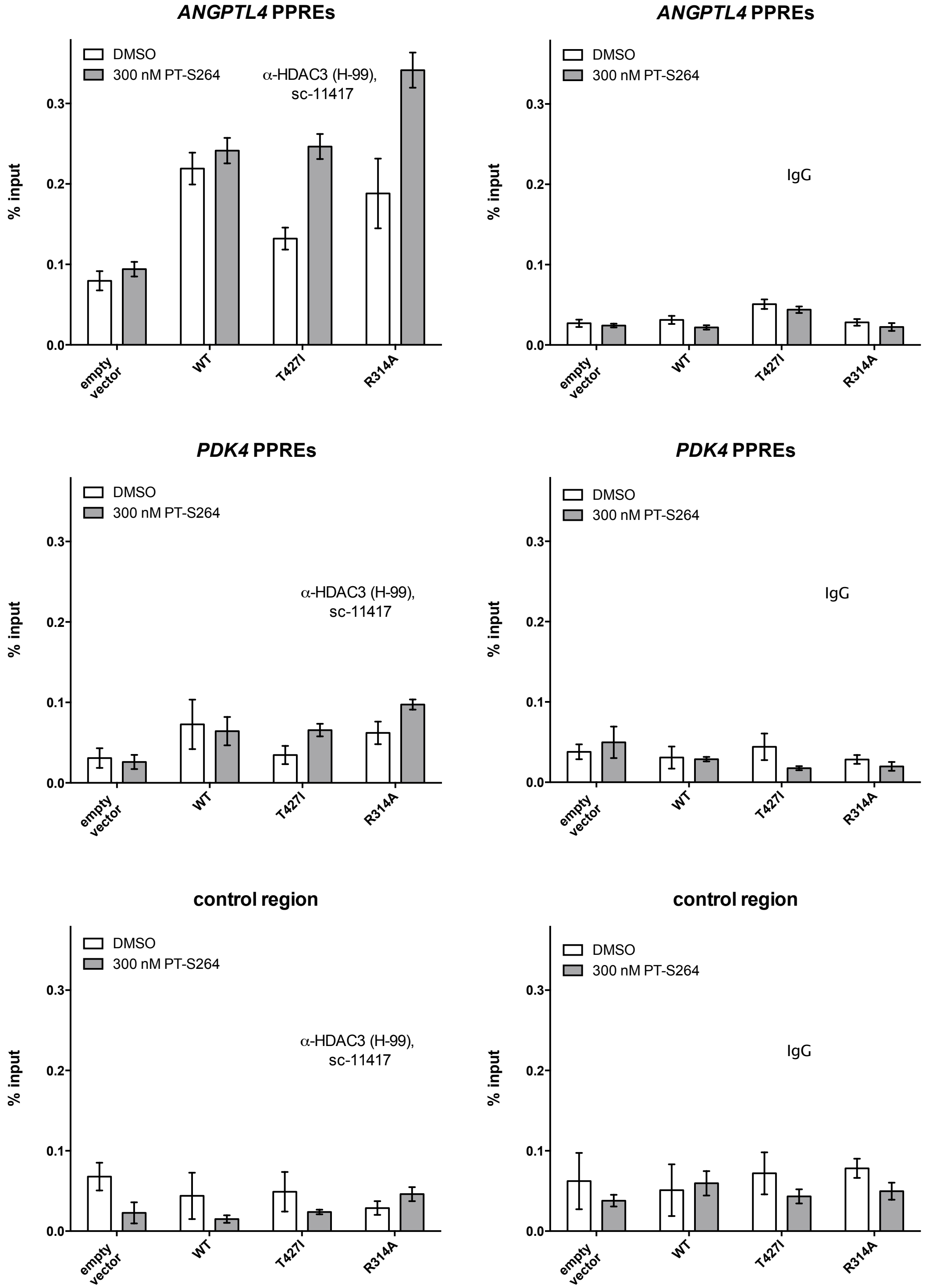


**Supp. fig. S10. HDAC3 recruitment to PPARβ/δ binding sites in cells expressing mutants deficient in basal repression.** 2B3 *PPARD* KO cells expressing the murine ecotropic receptor were retrovirally transduced with the empty vector or vectors carrying *PPARD* cDNAs as indicated. After selection, cells were treated with DMSO or 300 nM PT-S264 for 30 min and subjected to ChIP-qPCR analyses using an antibody against HDAC3 or an unspecific IgG pool. A representative experiment is shown. The error bars indicate standard deviation of three technical replicates.

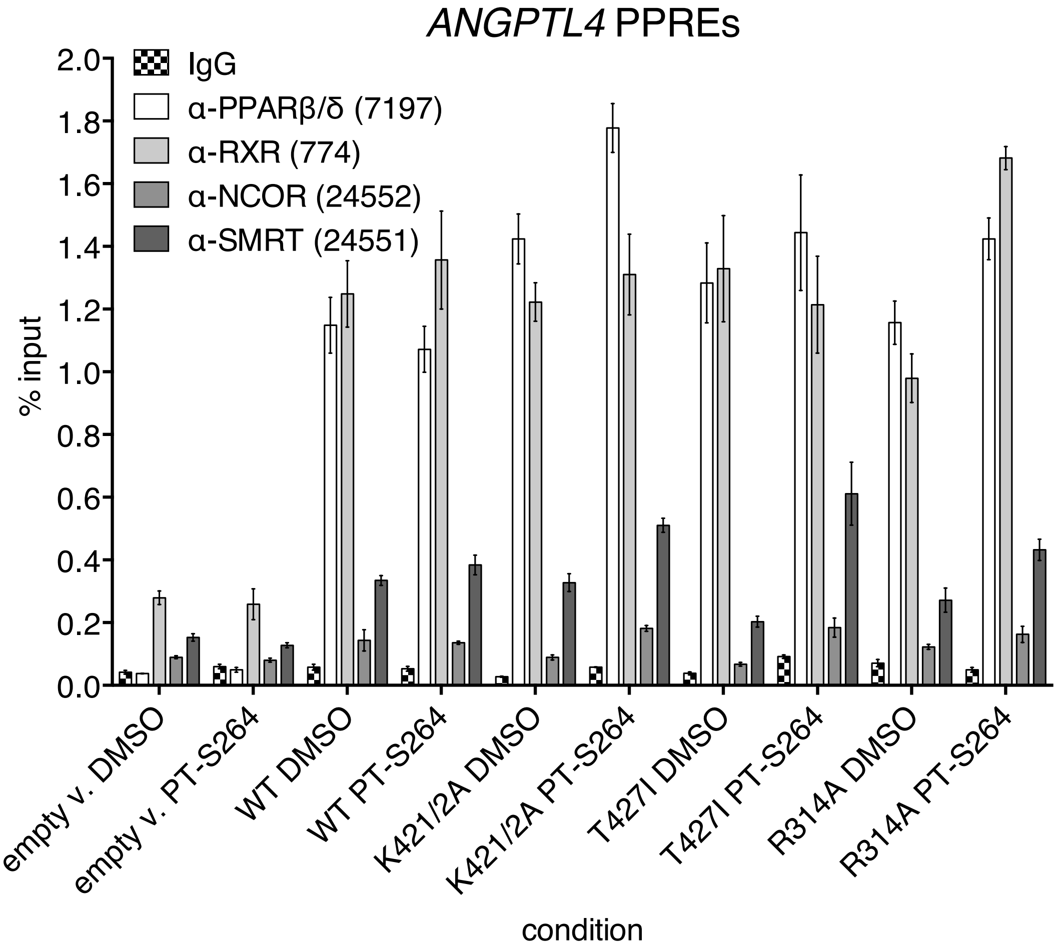


**Supp. fig. S11. PPARβ/δ, RXR, and NCOR recruitment to the PPAR-responsive enhancer of *ANGPTL4*.** 2B3 *PPARD* KO cells expressing the murine ecotropic receptor were retrovirally transduced with the empty vector or vectors carrying *PPARD* cDNAs as indicated. After selection, cells were treated with DMSO or 300 nM PT-S264 for 30 min and subjected to ChIP-qPCR analyses using antibodies against PPARβ/δ, RXR, NCOR, or an unspecific IgG pool. The numbers indicate the catalogue numbers of the antibodies. The primer pair amplifies the PPRE region of the *ANGPTL4* locus. The error bars indicate standard deviation of three technical replicates.

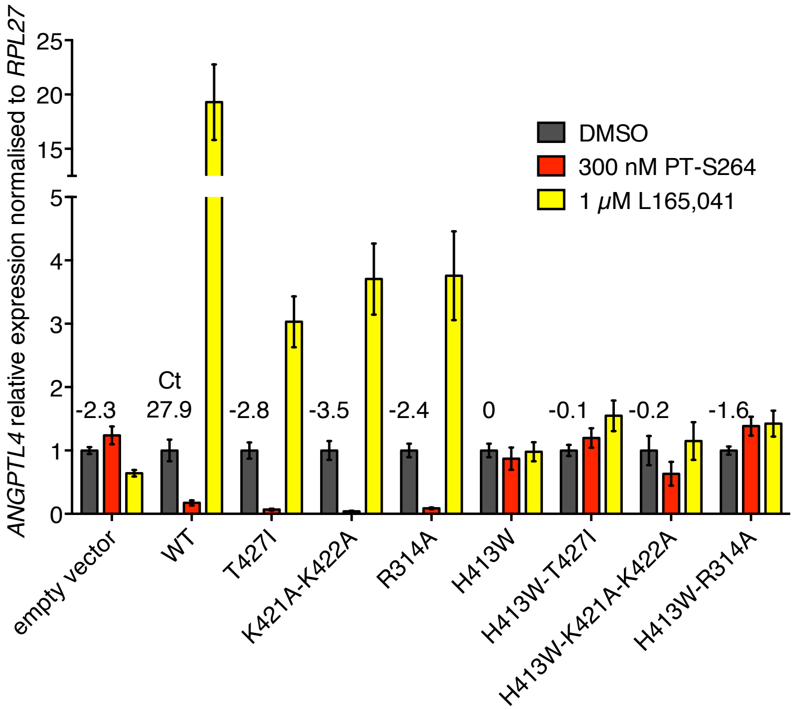


**Supp. fig. S12. Introduction of the H413W mutation abrogates ligand function in the T427I, K421A-K422A, and R314A mutants.** 2B3 *PPARD* KO cells expressing the murine ecotropic receptor were retrovirally transduced with the empty vector or vectors carrying *PPARD* cDNAs as indicated. The H413 amino acid residue faces inside the ligand binding pocket of PPARβ/δ, and mutation to tryptophane presumably causes steric hindrance of ligand binding. After selection, cells were treated with DMSO, 300 nM PT-S264, or 1 μM L165,041 for 6 h. *ANGPTL4* expression was measured by RT-qPCR using the *RPL27* transcript for normalisation. The error bars represent standard deviations calculated from the analyses of three technical replicates. The Ct value measured for the *ANGPTL4* transcript in the wildtype PPARβ/δ-reconstituted cells and relative changes for all other mutants are indicated.

**Supp. table 1.** RT-qPCR primers

| name | forward | reverse |
|---|---|---|
| *ANGPTL4* | TTTTGGTGAACTGCAAGATGA | GAAGTCCACTGAGCCATCGT |
| *PDK4* | TTATACATACTCCACTGCACCA | ATAGACTCAGAAGACAAAGCCT |
| *PLIN2* | GGAATCTTTAGATGACGTGATGG | CAAGTCTATGGTGGTGAAATCAA |
| *RPL27* | AAAGCTGTCATCGTGAAGAAC | GCTGTCACTTTGCGGGGGTAG |
| *TSC22D3* | CTTCTCTTCTCTGCTTGGAGGG | CGATCTTGTTGTCTATGGCCAC |

**Supp. table 2.** ChIP-qPCR primers

| name | forward | reverse |
|---|---|---|
| *ABCA1* TSS | GAGAACCGGCTCTGTTGGT | AATTGCGAGCGAGAGTGAGT |
| *ANGPTL4* –700 | AGGCAAGGACTTTTGGTGAG | GGAAGGAGGGAAAGAAAGG |
| *ANGPTL4* –500 | CTTTTTCCGTCCTTCCTTCC | GCGACAGAGCCAGAGTACG |
| *ANGPTL4* –275 | CGGGCTGGTCTGGAAGTC | GCCCGCCTCTAGTGTGAA |
| *ANGPTL4* TSS | TCCGCACCCAACTTATAAAAAC | GGATCACAGTCGTGTGAGGAT |
| *ANGPTL4* +250 | CCTAAGAGGATGAGCGGTGCT | TCGTCCCAGGACGCAAAG |
| *ANGPTL4* +300 | CACCGCCGTGCTACTGAG | GCCAGGACATTCATCTCGTC |
| *ANGPTL4* +500 | TCCACCGACCTCCCGTTAG | AACCAGCCCTGGGGACAC |
| *ANGPTL4* +750 | AGGAATTCAAGACCACCTAAAGC | GTGTGTGTGTGTGTAGAAGAGACG |
| *ANGPTL4* +2,000 | CGAATTCAGCATCTGCAAAG | GAGGGACCTTTTCTCCCTTG |
| *ANGPTL4* PPREs (+3,500) | CCCAGAGTGACCAGGAAGAC | CCTTACTGGATGGGAGGAAAG |
| *ANGPTL4* +5,000 | TAGGCAGATGGCAGAGAGGT | ACAGTGGATGACCAGGGAAG |
| *ANGPTL4* +7,000 | ACGATGGCTCAGTGGACTTC | CTGCCTCTGTCCCCACTAGA |
| *ANGPTL4* +10,500 | TGGTGCTGTTGTGTGTAGGTC | GCTTTTATTCCAAGAACTCTGTGAG |
| *PDK4* PPREs (–12,200) | GCAGAGTCAACAAGGGGAAG | ACTAGATGCCTGGGAGCTGA |
| *PDK4* TSS | GTCCCAAACAGGAGGAGTCA | CGGAGCCCATAGTTCTTTCTC |
| *PDK4* –4,500 (control) | GTATGTGTACTGGGGGGAC | CAGATGGCTCTTTTCGTTCC |
| *PLIN2* PPRE (–34,300) | CCAGTAAGCCTCTTAGCACCAC | TGCAACCTTTCACTTTGTGC |
| *PLIN2* TSS | GACGGACTGCAGCGAAAG | GCGAGGCGGGGTTTATAG |
| *TSC22D3* TSS | GCTGGAGTTGAAGGGAAGTG | AGGAGCCAAAATATCTCCGAGT |

**Supp. table 3.** Effects of the 80 mutations and the empty vector on basal and PPARβ/δ ligand-dependent *ANGPTL4* expression (165=L165,041; 264=PT-S264; dot means little or no effect, – means loss of effect, + means gain of effect relative to cells expressing the wildtype cDNA). AF: activating function; DBD: DNA binding domain; ZF: zinc finger; RFGR: R144-F145-G146-R147; Hx: helix x; S: β-pleated sheet, L-box: loop box, LXXLL: leucine-rich motif.

| mutant | position or del. | b.repr. | 264 | 165 | mutant | position or del. | b.repr. | 264 | 165 |
|---|---|---|---|---|---|---|---|---|---|
| 72–441 | ΔAF1 | – | • | – | M416K | H10-H11 | • | • | • |
| C91AE92A | DBD, ZF | – | – | – | M416R | H10-H11 | + | • | • |
| G146A | hinge, RFGR | – | – | – | M417A | H10-H11 | • | • | • |
| R147K | hinge, RFGR | – | • | – | I420A | H10-H11 | • | • | • |
| R154K | hinge | • | • | • | I420R | H10-H11 | • | – | – |
| K155R | hinge | • | • | • | K421A | H10-H11 | – | + | – |
| K229R | H2'-H3 loop | • | • | • | K421R | H10-H11 | – | • | • |
| K239R | H3 | • | • | • | K422A | H10-H11 | • | • | • |
| E240A | H3 | • | • | • | K422R | H10-H11 | • | • | • |
| V254A | H3 | + | – | • | K421A-K422A | H10-H11 | – | + | – |
| V254R | H3 | • | • | – | K421R-K422R | H10-H11 | – | • | • |
| V257R | H3 | – | – | – | T423A | L-box | + | – | • |
| T261R | H3 | – | – | – | T423D | L-box | • | • | • |
| V279R | H4 | – | – | – | E424A | L-box | • | • | • |
| T261R-V279R | H3,H4 | – | – | – | T425A | L-box | • | • | • |
| L281A | H4 | • | • | • | T425D | L-box | • | • | • |
| K283A | H4 | • | • | – | E426A | L-box | • | • | • |
| K300A | S2 | • | • | • | T427A | L-box | • | • | • |
| K300R | S2 | • | • | • | T427D | L-box | + | • | • |
| R314A | H6 | – | • | – | T427F | L-box | – | • | • |
| R314K | H6 | • | • | • | T427I | L-box | – | + | – |
| E315A | H6 | • | • | • | T423A-T427A | L-box | • | • | • |
| K318A | H6 | • | • | • | T423A-T427D | L-box | • | • | – |
| K318R | H6 | • | • | • | T423D-T427A | L-box | • | • | • |
| R321A | H6-H7 loop | + | – | • | T423D-T427D | L-box | • | – | • |
| R321K | H6-H7 loop | • | – | – | S428A | L-box | • | • | • |
| K322A | H6-H7 loop | • | • | • | S428D | L-box | • | • | • |
| K322R | H6-H7 loop | • | • | • | L429A | LXXLL, H12 | • | – | – |
| F324A | H6-H7 loop | • | – | – | H430K | LXXLL, H12 | • | • | – |
| D326A | H7 | • | • | • | L432A | LXXLL, H12 | + | – | – |
| I327A | H7 | + | – | • | L432I | LXXLL, H12 | • | – | • |
| I328A | H7 | + | • | • | L433A | LXXLL, H12 | • | – | – |
| E329A | H8 | – | • | – | L433F | LXXLL, H12 | + | – | – |
| E333A | H8 | • | • | • | L433I | LXXLL, H12 | + | – | – |
| A404K | H10-H11 | – | – | – | E435A | H12 (AF2) | • | • | – |
| A404W | H10-H11 | – | – | – | Y437A | H12 (AF2) | • | – | – |
| L406R | H10-H11 | – | – | – | Y437F | H12 (AF2) | • | • | • |
| H413A | H10-H11 | • | • | – | D439A | H12 (AF2) | – | – | – |
| H413W | H10-H11 | • | – | – | 1–421 | ΔL-box, ΔAF2 | • | • | – |
| M416A | H10-H11 | • | • | • | 1–428 | ΔAF2 | • | • | – |
| | | | | | empty vector | Δall | – | – | – |

Appendix A. Expression data for the PPARβ/δ mutant screen, sorted by positions.

| mutant | position | basal repression | | PT-S264 | | | L165,041 | | | n |
|---|---|---|---|---|---|---|---|---|---|---|
| | | basal Ct | change vs WT | Mean | SD | change vs WT | MW | SD | change vs WT | |
| WT (Mean) | | 28.9 | 0.0 | 0.183 | 0.052 | 0.00 | 19.840 | 6.979 | 0.00 | 20 |
| 72-441 | ΔAF1 | 27.0 | -1.9 | 0.379 | 0.105 | -1.05 | 6.586 | 3.113 | -1.59 | 3 |
| C91A-E92A | DBD | 27.3 | -1.6 | 1.020 | 0.194 | -2.48 | 1.220 | 0.173 | -4.02 | 8 |
| G146A | RFGR | 26.0 | -2.9 | 0.877 | 0.090 | -2.26 | 1.266 | 0.060 | -3.97 | 1 |
| R147K | RFGR | 27.4 | -1.5 | 0.184 | 0.034 | -0.01 | 6.869 | 0.870 | -1.53 | 1 |
| R154K | Hinge | 28.0 | -0.9 | 0.240 | 0.051 | -0.39 | 16.000 | 1.711 | -0.31 | 1 |
| K155R | Hinge | 28.4 | -0.5 | 0.193 | 0.098 | -0.08 | 20.393 | 2.208 | 0.04 | 1 |
| K229R | between H2'/H3 | 28.5 | -0.4 | 0.291 | 0.034 | -0.67 | 19.027 | 1.684 | -0.06 | 1 |
| K239R | H3 | 29.6 | 0.7 | 0.283 | 0.084 | -0.63 | 28.247 | 3.346 | 0.51 | 1 |
| E240A | H3 | 30.1 | 1.2 | 0.363 | 0.085 | -0.99 | 16.111 | 3.019 | -0.30 | 1 |
| V254A | H3 | 30.4 | 1.5 | 0.651 | 0.225 | -1.83 | 24.062 | 7.498 | 0.28 | 2 |
| V254R | H3 | 29.6 | 0.7 | 0.464 | 0.106 | -1.34 | 1.508 | 0.151 | -3.72 | 2 |
| V257R | H3 | 27.0 | -1.9 | 1.012 | 0.142 | -2.47 | 1.250 | 0.260 | -3.99 | 3 |
| T261R | H3 | 26.8 | -2.1 | 0.982 | 0.136 | -2.42 | 1.366 | 0.179 | -3.86 | 3 |
| V279R | H4 | 26.8 | -2.1 | 0.650 | 0.030 | -1.83 | 1.128 | 0.077 | -4.14 | 3 |
| T261R-V279R | H3/H4 | 26.7 | -2.2 | 0.898 | 0.016 | -2.29 | 1.035 | 0.007 | -4.26 | 2 |
| L281A | H4 | 28.2 | -0.7 | 0.503 | 0.090 | -1.46 | 11.158 | 1.335 | -0.83 | 1 |
| K283A | H4 | 28.7 | -0.2 | 0.516 | 0.059 | -1.50 | 2.782 | 0.823 | -2.83 | 2 |
| K300A | S2 | 29.1 | 0.2 | 0.295 | 0.094 | -0.69 | 24.930 | 2.444 | 0.33 | 1 |
| K300R | S2 | 29.0 | 0.1 | 0.149 | 0.083 | 0.30 | 23.918 | 2.663 | 0.27 | 1 |
| R314A | H6 | 26.5 | -2.4 | 0.083 | 0.031 | 1.14 | 5.084 | 1.219 | -1.96 | 5 |
| R314K | H6 | 28.8 | -0.1 | 0.163 | 0.033 | 0.17 | 14.221 | 1.954 | -0.48 | 1 |
| E315A | H6 | 29.1 | 0.2 | 0.204 | 0.042 | -0.16 | 17.268 | 1.795 | -0.20 | 1 |
| K318A | H6 | 28.8 | -0.1 | 0.199 | 0.044 | -0.12 | 13.830 | 2.605 | -0.52 | 1 |
| K318R | H6 | 29.2 | 0.3 | 0.124 | 0.018 | 0.56 | 25.634 | 4.667 | 0.37 | 1 |
| R321A | between H6/H7 | 30.8 | 1.9 | 0.547 | 0.061 | -1.58 | 33.591 | 3.374 | 0.76 | 1 |
| R321K | between. H6/H7 | 28.1 | -0.8 | 0.780 | 0.023 | -2.09 | 7.312 | 2.103 | -1.44 | 3 |
| K322A | between H6/H7 | 28.0 | -0.9 | 0.186 | 0.034 | -0.02 | 18.379 | 2.493 | -0.11 | 1 |
| K322R | between. H6/H7 | 28.4 | -0.5 | 0.186 | 0.038 | -0.02 | 12.295 | 1.734 | -0.69 | 1 |
| F324A | between H6/H7 | 28.1 | -0.8 | 0.540 | 0.183 | -1.56 | 2.269 | 0.198 | -3.13 | 3 |
| D326A | H7 | 30.3 | 1.4 | 0.454 | 0.092 | -1.31 | 49.180 | 7.973 | 1.31 | 1 |
| I327A | H7 | 31.6 | 2.7 | 0.675 | 0.201 | -1.88 | 7.950 | 3.227 | -1.32 | 3 |
| I328A | H7 | 30.7 | 1.8 | 0.406 | 0.028 | -1.15 | 26.355 | 1.582 | 0.41 | 1 |
| E329A | H7 | 27.2 | -1.7 | 0.207 | 0.038 | -0.18 | 6.277 | 0.847 | -1.66 | 1 |
| E333A | H7 | 28.7 | -0.2 | 0.136 | 0.067 | 0.43 | 10.411 | 1.653 | -0.93 | 1 |
| A404K | H10/11 | 27.1 | -1.8 | 0.979 | 0.104 | -2.42 | 2.346 | 0.263 | -3.08 | 1 |
| A404W | H10/11 | 27.4 | -1.5 | 0.774 | 0.086 | -2.08 | 2.549 | 0.252 | -2.96 | 1 |
| L406R | H10/11 | 26.5 | -2.4 | 1.057 | 0.111 | -2.53 | 1.206 | 0.124 | -4.04 | 1 |
| H413A | H10/11 | 28.1 | -0.8 | 0.454 | 0.088 | -1.31 | 3.294 | 0.381 | -2.59 | 1 |
| H413W | H10/11 | 28.5 | -0.4 | 0.920 | 0.090 | -2.33 | 1.196 | 0.189 | -4.05 | 5 |
| M416A | H10/11 | 28.9 | 0.0 | 0.358 | 0.070 | -0.97 | 30.484 | 2.966 | 0.62 | 1 |
| M416K | H10/11 | 28.0 | -0.9 | 0.268 | 0.040 | -0.55 | 15.455 | 1.913 | -0.36 | 1 |
| M416R | H10/11 | 30.5 | 1.6 | 0.218 | 0.056 | -0.25 | 22.943 | 4.186 | 0.21 | 1 |
| M417A | H10/11 | 29.4 | 0.5 | 0.515 | 0.083 | -1.49 | 12.405 | 2.419 | -0.68 | 2 |
| I420A | H10/11 | 29.5 | 0.6 | 0.354 | 0.118 | -0.95 | 13.269 | 2.499 | -0.58 | 1 |
| I420R | H10/11 | 28.0 | -0.9 | 0.655 | 0.105 | -1.84 | 5.205 | 1.049 | -1.93 | 1 |
| K421A | H10/11 | 24.9 | -4.0 | 0.043 | 0.004 | 2.09 | 4.412 | 1.176 | -2.17 | 3 |
| K421R | H10/11 | 26.9 | -2.0 | 0.095 | 0.005 | 0.95 | 7.632 | 1.816 | -1.38 | 2 |
| K422A | H10/11 | 28.7 | -0.2 | 0.133 | 0.013 | 0.46 | 32.728 | 4.286 | 0.72 | 2 |
| K422R | H10/11 | 28.0 | -0.9 | 0.182 | 0.066 | 0.01 | 13.455 | 0.093 | -0.56 | 2 |
| K421A-K422A | H10/11 | 26.4 | -2.5 | 0.037 | 0.009 | 2.31 | 3.346 | 0.264 | -2.57 | 6 |
| K421R-K422R | H10/11 | 27.4 | -1.5 | 0.104 | 0.025 | 0.82 | 13.107 | 0.726 | -0.60 | 2 |
| T423A | L-box | 31.1 | 2.2 | 0.678 | 0.049 | -1.89 | 43.518 | 9.927 | 1.13 | 2 |
| T423D | L-box | 29.2 | 0.3 | 0.257 | 0.042 | -0.49 | 30.484 | 4.059 | 0.62 | 1 |
| E424A | L-box | 28.8 | -0.1 | 0.262 | 0.084 | -0.52 | 15.137 | 3.025 | -0.39 | 1 |
| T425A | L-box | 29.9 | 1.0 | 0.426 | 0.060 | -1.22 | 44.632 | 5.241 | 1.17 | 1 |
| T425D | L-box | 30.1 | 1.2 | 0.374 | 0.075 | -1.03 | 41.355 | 5.207 | 1.06 | 1 |
| E426A | L-box | 27.9 | -1.0 | 0.306 | 0.065 | -0.74 | 15.560 | 1.320 | -0.35 | 1 |
| T427A | L-box | 28.2 | -0.7 | 0.140 | 0.038 | 0.39 | 15.137 | 1.517 | -0.39 | 1 |
| T427F | L-box | 26.6 | -2.3 | 0.112 | 0.028 | 0.71 | 9.189 | 1.515 | -1.11 | 2 |
| T427I | L-box | 26.2 | -2.7 | 0.037 | 0.018 | 2.31 | 3.307 | 0.561 | -2.58 | 8 |
| T427D | L-box | 31.5 | 2.6 | 0.465 | 0.138 | -1.35 | 15.263 | 5.367 | -0.38 | 5 |
| T423A-T427A | L-box | 29.8 | 0.9 | 0.415 | 0.172 | -1.18 | 48.961 | 18.688 | 1.30 | 2 |
| T423D-T427D | L-box | 30.2 | 1.3 | 0.840 | 0.227 | -2.20 | 18.292 | 5.094 | -0.12 | 3 |
| T423A-T427D | L-box | 29.4 | 0.5 | 0.505 | 0.090 | -1.46 | 7.010 | 0.990 | -1.50 | 2 |
| T423D-T427A | L-box | 28.8 | -0.1 | 0.230 | 0.039 | -0.33 | 28.791 | 0.989 | 0.54 | 3 |
| S428A | L-box | 27.4 | -1.5 | 0.156 | 0.051 | 0.23 | 19.160 | 4.785 | -0.05 | 1 |
| S428D | L-box | 29.3 | 0.4 | 0.342 | 0.054 | -0.90 | 36.002 | 5.205 | 0.86 | 1 |
| L429A | LXXLL | 29.9 | 1.0 | 0.926 | 0.424 | -2.34 | 2.904 | 1.947 | -2.77 | 3 |
| H430K | LXXLL | 30.1 | 1.2 | 0.271 | 0.126 | -0.57 | 2.770 | 0.444 | -2.84 | 1 |
| L432A | LXXLL, H12 | 30.8 | 1.9 | 1.020 | 0.392 | -2.48 | 2.904 | 0.147 | -2.77 | 3 |
| L432I | LXXLL, H12 | 30.3 | 1.4 | 0.525 | 0.229 | -1.52 | 19.562 | 6.226 | -0.02 | 1 |
| L433A | LXXLL, H12 | 29.4 | 0.5 | 0.532 | 0.161 | -1.54 | 1.283 | 0.287 | -3.95 | 1 |
| L433F | LXXLL, H12 | 31.0 | 2.1 | 0.708 | 0.164 | -1.95 | 1.382 | 0.204 | -3.84 | 3 |
| L433I | LXXLL, H12 | 30.9 | 2.0 | 0.742 | 0.054 | -2.02 | 31.651 | 24.418 | 0.67 | 3 |
| E435A | H12 | 28.9 | 0.0 | 0.340 | 0.026 | -0.89 | 4.628 | 0.960 | -2.10 | 2 |
| Y437A | H12 | 29.5 | 0.6 | 0.591 | 0.050 | -1.69 | 1.461 | 0.234 | -3.76 | 3 |
| Y437F | H12 | 28.1 | -0.8 | 0.137 | 0.025 | 0.42 | 8.694 | 1.134 | -1.190 | 1 |
| D439A | H12 | 26.3 | -2.6 | 0.836 | 0.120 | -2.19 | 1.486 | 0.360 | -3.74 | 3 |
| 1-421 | ΔL-box+AF2 | 28.6 | -0.3 | 0.382 | 0.044 | -1.06 | 1.233 | 0.068 | -4.01 | 3 |
| 1-428 | ΔAF2 | 28.5 | -0.4 | 0.426 | 0.079 | -1.22 | 1.040 | 0.199 | -4.25 | 3 |
| empty vector | | 26.3 | -2.6 | 0.993 | 0.116 | -2.44 | 1.273 | 0.234 | -3.96 | 9 |

| |
|---|
| wildtype |
| n > 1 |
| change ≤ -1,5 |
| change ≥ 1,5 |

Appendix B. Expression data for the PPARβ/δ mutant screen, sorted by effect on basal expression.

| mutant | position | basal repression | | PT-S264 | | | L165,041 | | | n |
|---|---|---|---|---|---|---|---|---|---|---|
| | | basal Ct | change vs WT | Mean | SD | change vs WT | MW | SD | change vs WT | |
| WT (Mean) | | 28.9 | 0.0 | 0.183 | 0.052 | 0.00 | 19.840 | 6.979 | 0.00 | 20 |
| K421A | H10/11 | 24.9 | -4.0 | 0.043 | 0.004 | 2.09 | 4.412 | 1.176 | -2.17 | 3 |
| G146A | RFGR | 26.0 | -2.9 | 0.877 | 0.090 | -2.26 | 1.266 | 0.060 | -3.97 | 1 |
| T427I | L-box | 26.2 | -2.7 | 0.037 | 0.018 | 2.31 | 3.307 | 0.561 | -2.58 | 8 |
| D439A | H12 | 26.3 | -2.6 | 0.836 | 0.120 | -2.19 | 1.486 | 0.360 | -3.74 | 3 |
| empty vector | | 26.3 | -2.6 | 0.993 | 0.116 | -2.44 | 1.273 | 0.234 | -3.96 | 9 |
| K421A-K422A | H10/11 | 26.4 | -2.5 | 0.037 | 0.009 | 2.31 | 3.346 | 0.264 | -2.57 | 6 |
| R314A | H6 | 26.5 | -2.4 | 0.083 | 0.031 | 1.14 | 5.084 | 1.219 | -1.96 | 5 |
| L406R | H10/11 | 26.5 | -2.4 | 1.057 | 0.111 | -2.53 | 1.206 | 0.124 | -4.04 | 1 |
| T427F | L-box | 26.6 | -2.3 | 0.112 | 0.028 | 0.71 | 9.189 | 1.515 | -1.11 | 2 |
| T261R-V279R | H3/H4 | 26.7 | -2.2 | 0.898 | 0.016 | -2.29 | 1.035 | 0.007 | -4.26 | 2 |
| T261R | H3 | 26.8 | -2.1 | 0.982 | 0.136 | -2.42 | 1.366 | 0.179 | -3.86 | 3 |
| V279R | H4 | 26.8 | -2.1 | 0.650 | 0.030 | -1.83 | 1.128 | 0.077 | -4.14 | 3 |
| K421R | H10/11 | 26.9 | -2.0 | 0.095 | 0.005 | 0.95 | 7.632 | 1.816 | -1.38 | 2 |
| 72-441 | ΔAF1 | 27.0 | -1.9 | 0.379 | 0.105 | -1.05 | 6.586 | 3.113 | -1.59 | 3 |
| V257R | H3 | 27.0 | -1.9 | 1.012 | 0.142 | -2.47 | 1.250 | 0.260 | -3.99 | 3 |
| A404K | H10/11 | 27.1 | -1.8 | 0.979 | 0.104 | -2.42 | 2.346 | 0.263 | -3.08 | 1 |
| E329A | H7 | 27.2 | -1.7 | 0.207 | 0.038 | -0.18 | 6.277 | 0.847 | -1.66 | 1 |
| C91A-E92A | DBD | 27.3 | -1.6 | 1.020 | 0.194 | -2.48 | 1.220 | 0.173 | -4.02 | 8 |
| R147K | RFGR | 27.4 | -1.5 | 0.184 | 0.034 | -0.01 | 6.869 | 0.870 | -1.53 | 1 |
| A404W | H10/11 | 27.4 | -1.5 | 0.774 | 0.086 | -2.08 | 2.549 | 0.252 | -2.96 | 1 |
| K421R-K422R | H10/11 | 27.4 | -1.5 | 0.104 | 0.025 | 0.82 | 13.107 | 0.726 | -0.60 | 2 |
| S428A | L-box | 27.4 | -1.5 | 0.156 | 0.051 | 0.23 | 19.160 | 4.785 | -0.05 | 1 |
| E426A | L-box | 27.9 | -1.0 | 0.306 | 0.065 | -0.74 | 15.560 | 1.320 | -0.35 | 1 |
| R154K | Hinge | 28.0 | -0.9 | 0.240 | 0.051 | -0.39 | 16.000 | 1.711 | -0.31 | 1 |
| K322A | between H6/H7 | 28.0 | -0.9 | 0.186 | 0.034 | -0.02 | 18.379 | 2.493 | -0.11 | 1 |
| M416K | H10/11 | 28.0 | -0.9 | 0.268 | 0.040 | -0.55 | 15.455 | 1.913 | -0.36 | 1 |
| I420R | H10/11 | 28.0 | -0.9 | 0.655 | 0.105 | -1.84 | 5.205 | 1.049 | -1.93 | 1 |
| K422R | H10/11 | 28.0 | -0.9 | 0.182 | 0.066 | 0.01 | 13.455 | 0.093 | -0.56 | 2 |
| R321K | between. H6/H7 | 28.1 | -0.8 | 0.780 | 0.023 | -2.09 | 7.312 | 2.103 | -1.44 | 3 |
| F324A | between H6/H7 | 28.1 | -0.8 | 0.540 | 0.183 | -1.56 | 2.269 | 0.198 | -3.13 | 3 |
| H413A | H10/11 | 28.1 | -0.8 | 0.454 | 0.088 | -1.31 | 3.294 | 0.381 | -2.59 | 1 |
| Y437F | H12 | 28.1 | -0.8 | 0.137 | 0.025 | 0.42 | 8.694 | 1.134 | -1.19 | 1 |
| L281A | H4 | 28.2 | -0.7 | 0.503 | 0.090 | -1.46 | 11.158 | 1.335 | -0.83 | 1 |
| T427A | L-box | 28.2 | -0.7 | 0.140 | 0.038 | 0.39 | 15.137 | 1.517 | -0.39 | 1 |
| K155R | Hinge | 28.4 | -0.5 | 0.193 | 0.098 | -0.08 | 20.393 | 2.208 | 0.04 | 1 |
| K322R | between. H6/H7 | 28.4 | -0.5 | 0.186 | 0.038 | -0.02 | 12.295 | 1.734 | -0.69 | 1 |
| K229R | between H2'/H3 | 28.5 | -0.4 | 0.291 | 0.034 | -0.67 | 19.027 | 1.684 | -0.06 | 1 |
| H413W | H10/11 | 28.5 | -0.4 | 0.920 | 0.090 | -2.33 | 1.196 | 0.189 | -4.05 | 5 |
| 1-428 | ΔAF2 | 28.5 | -0.4 | 0.426 | 0.079 | -1.22 | 1.040 | 0.199 | -4.25 | 3 |
| 1-421 | ΔL-box+AF2 | 28.6 | -0.3 | 0.382 | 0.044 | -1.06 | 1.233 | 0.068 | -4.01 | 3 |
| K283A | H4 | 28.7 | -0.2 | 0.516 | 0.059 | -1.50 | 2.782 | 0.823 | -2.83 | 2 |
| E333A | H7 | 28.7 | -0.2 | 0.136 | 0.067 | 0.43 | 10.411 | 1.653 | -0.93 | 1 |
| K422A | H10/11 | 28.7 | -0.2 | 0.133 | 0.013 | 0.46 | 32.728 | 4.286 | 0.72 | 2 |
| R314K | H6 | 28.8 | -0.1 | 0.163 | 0.033 | 0.17 | 14.221 | 1.954 | -0.48 | 1 |
| K318A | H6 | 28.8 | -0.1 | 0.199 | 0.044 | -0.12 | 13.830 | 2.605 | -0.52 | 1 |
| E424A | L-box | 28.8 | -0.1 | 0.262 | 0.084 | -0.52 | 15.137 | 3.025 | -0.39 | 1 |
| T423D-T427A | L-box | 28.8 | -0.1 | 0.230 | 0.039 | -0.33 | 28.791 | 0.989 | 0.54 | 3 |
| M416A | H10/11 | 28.9 | 0.0 | 0.358 | 0.070 | -0.97 | 30.484 | 2.966 | 0.62 | 1 |
| E435A | H12 | 28.9 | 0.0 | 0.340 | 0.026 | -0.89 | 4.628 | 0.960 | -2.10 | 2 |
| WT (Mean) | | 28.9 | 0.0 | 0.183 | 0.052 | 0.00 | 19.840 | 6.979 | 0.00 | 20 |
| K300R | S2 | 29.0 | 0.1 | 0.149 | 0.083 | 0.30 | 23.918 | 2.663 | 0.27 | 1 |
| K300A | S2 | 29.1 | 0.2 | 0.295 | 0.094 | -0.69 | 24.930 | 2.444 | 0.33 | 1 |
| E315A | H6 | 29.1 | 0.2 | 0.204 | 0.042 | -0.16 | 17.268 | 1.795 | -0.20 | 1 |
| K318R | H6 | 29.2 | 0.3 | 0.124 | 0.018 | 0.56 | 25.634 | 4.667 | 0.37 | 1 |
| T423D | L-box | 29.2 | 0.3 | 0.257 | 0.042 | -0.49 | 30.484 | 4.059 | 0.62 | 1 |
| S428D | L-box | 29.3 | 0.4 | 0.342 | 0.054 | -0.90 | 36.002 | 5.205 | 0.86 | 1 |
| M417A | H10/11 | 29.4 | 0.5 | 0.515 | 0.083 | -1.49 | 12.405 | 2.419 | -0.68 | 2 |
| T423A-T427D | L-box | 29.4 | 0.5 | 0.505 | 0.090 | -1.46 | 7.010 | 0.990 | -1.50 | 2 |
| L433A | LXXLL, H12 | 29.4 | 0.5 | 0.532 | 0.161 | -1.54 | 1.283 | 0.287 | -3.95 | 1 |
| I420A | H10/11 | 29.5 | 0.6 | 0.354 | 0.118 | -0.95 | 13.269 | 2.499 | -0.58 | 1 |
| Y437A | H12 | 29.5 | 0.6 | 0.591 | 0.050 | -1.69 | 1.461 | 0.234 | -3.76 | 3 |
| K239R | H3 | 29.6 | 0.7 | 0.283 | 0.084 | -0.63 | 28.247 | 3.346 | 0.51 | 1 |
| V254R | H3 | 29.6 | 0.7 | 0.464 | 0.106 | -1.34 | 1.508 | 0.151 | -3.72 | 2 |
| T423A-T427A | L-box | 29.8 | 0.9 | 0.415 | 0.172 | -1.18 | 48.961 | 18.688 | 1.30 | 2 |
| T425A | L-box | 29.9 | 1.0 | 0.426 | 0.060 | -1.22 | 44.632 | 5.241 | 1.17 | 1 |
| L429A | LXXLL | 29.9 | 1.0 | 0.926 | 0.424 | -2.34 | 2.904 | 1.947 | -2.77 | 3 |
| E240A | H3 | 30.1 | 1.2 | 0.363 | 0.085 | -0.99 | 16.111 | 3.019 | -0.30 | 1 |
| T425D | L-box | 30.1 | 1.2 | 0.374 | 0.075 | -1.03 | 41.355 | 5.207 | 1.06 | 1 |
| H430K | LXXLL | 30.1 | 1.2 | 0.271 | 0.126 | -0.57 | 2.770 | 0.444 | -2.84 | 1 |
| T423D-T427D | L-box | 30.2 | 1.3 | 0.840 | 0.227 | -2.20 | 18.292 | 5.094 | -0.12 | 3 |
| D326A | H7 | 30.3 | 1.4 | 0.454 | 0.092 | -1.31 | 49.180 | 7.973 | 1.31 | 1 |
| L432I | LXXLL, H12 | 30.3 | 1.4 | 0.525 | 0.229 | -1.52 | 19.562 | 6.226 | -0.02 | 1 |
| V254A | H3 | 30.4 | 1.5 | 0.651 | 0.225 | -1.83 | 24.062 | 7.498 | 0.28 | 2 |
| M416R | H10/11 | 30.5 | 1.6 | 0.218 | 0.056 | -0.25 | 22.943 | 4.186 | 0.21 | 1 |
| I328A | H7 | 30.7 | 1.8 | 0.406 | 0.028 | -1.15 | 26.355 | 1.582 | 0.41 | 1 |
| R321A | between H6/H7 | 30.8 | 1.9 | 0.547 | 0.061 | -1.58 | 33.591 | 3.374 | 0.76 | 1 |
| L432A | LXXLL, H12 | 30.8 | 1.9 | 1.020 | 0.392 | -2.48 | 2.904 | 0.147 | -2.77 | 3 |
| L433I | LXXLL, H12 | 30.9 | 2.0 | 0.742 | 0.054 | -2.02 | 31.651 | 24.418 | 0.67 | 3 |
| L433F | LXXLL, H12 | 31.0 | 2.1 | 0.708 | 0.164 | -1.95 | 1.382 | 0.204 | -3.84 | 3 |
| T423A | L-box | 31.1 | 2.2 | 0.678 | 0.049 | -1.89 | 43.518 | 9.927 | 1.13 | 2 |
| T427D | L-box | 31.5 | 2.6 | 0.465 | 0.138 | -1.35 | 15.263 | 5.367 | -0.38 | 5 |
| I327A | H7 | 31.6 | 2.7 | 0.675 | 0.201 | -1.88 | 7.950 | 3.227 | -1.32 | 3 |

| |
|---|
| wildtype |
| n > 1 |
| change ≤ -1,5 |
| change ≥ 1,5 |

Appendix C. Expression data for the PPARβ/δ mutant screen, sorted by effect on L165,041–dependent change.

| mutant | position | basal repression | | PT-S264 | | | L165,041 | | | n |
|---|---|---|---|---|---|---|---|---|---|---|
| | | basal Ct | change vs WT | Mean | SD | change vs WT | MW | SD | change vs WT | |
| WT (Mean) | | 28.9 | 0.0 | 0.183 | 0.052 | 0.00 | 19.840 | 6.979 | 0.00 | 20 |
| T261R-V279R | H3/H4 | 26.7 | -2.2 | 0.898 | 0.016 | -2.29 | 1.035 | 0.007 | -4.26 | 2 |
| 1-428 | ΔAF2 | 28.5 | -0.4 | 0.426 | 0.079 | -1.22 | 1.040 | 0.199 | -4.25 | 3 |
| V279R | H4 | 26.8 | -2.1 | 0.650 | 0.030 | -1.83 | 1.128 | 0.077 | -4.14 | 3 |
| H413W | H10/11 | 28.5 | -0.4 | 0.920 | 0.090 | -2.33 | 1.196 | 0.189 | -4.05 | 5 |
| L406R | H10/11 | 26.5 | -2.4 | 1.057 | 0.111 | -2.53 | 1.206 | 0.124 | -4.04 | 1 |
| C91A-E92A | DBD | 27.3 | -1.6 | 1.020 | 0.194 | -2.48 | 1.220 | 0.173 | -4.02 | 8 |
| 1-421 | ΔL-box+AF2 | 28.6 | -0.3 | 0.382 | 0.044 | -1.06 | 1.233 | 0.068 | -4.01 | 3 |
| V257R | H3 | 27.0 | -1.9 | 1.012 | 0.142 | -2.47 | 1.250 | 0.260 | -3.99 | 3 |
| G146A | RFGR | 26.0 | -2.9 | 0.877 | 0.090 | -2.26 | 1.266 | 0.060 | -3.97 | 1 |
| empty vector | | 26.3 | -2.6 | 0.993 | 0.116 | -2.44 | 1.273 | 0.234 | -3.96 | 9 |
| L433A | LXXLL, H12 | 29.4 | 0.5 | 0.532 | 0.161 | -1.54 | 1.283 | 0.287 | -3.95 | 1 |
| T261R | H3 | 26.8 | -2.1 | 0.982 | 0.136 | -2.42 | 1.366 | 0.179 | -3.86 | 3 |
| L433F | LXXLL, H12 | 31.0 | 2.1 | 0.708 | 0.164 | -1.95 | 1.382 | 0.204 | -3.84 | 3 |
| Y437A | H12 | 29.5 | 0.6 | 0.591 | 0.050 | -1.69 | 1.461 | 0.234 | -3.76 | 3 |
| D439A | H12 | 26.3 | -2.6 | 0.836 | 0.120 | -2.19 | 1.486 | 0.360 | -3.74 | 3 |
| V254R | H3 | 29.6 | 0.7 | 0.464 | 0.106 | -1.34 | 1.508 | 0.151 | -3.72 | 2 |
| F324A | between H6/H7 | 28.1 | -0.8 | 0.540 | 0.183 | -1.56 | 2.269 | 0.198 | -3.13 | 3 |
| A404K | H10/11 | 27.1 | -1.8 | 0.979 | 0.104 | -2.42 | 2.346 | 0.263 | -3.08 | 1 |
| A404W | H10/11 | 27.4 | -1.5 | 0.774 | 0.086 | -2.08 | 2.549 | 0.252 | -2.96 | 1 |
| H430K | LXXLL | 30.1 | 1.2 | 0.271 | 0.126 | -0.57 | 2.770 | 0.444 | -2.84 | 1 |
| K283A | H4 | 28.7 | -0.2 | 0.516 | 0.059 | -1.50 | 2.782 | 0.823 | -2.83 | 2 |
| L432A | LXXLL, H12 | 30.8 | 1.9 | 1.020 | 0.392 | -2.48 | 2.904 | 0.147 | -2.77 | 3 |
| L429A | LXXLL | 29.9 | 1.0 | 0.926 | 0.424 | -2.34 | 2.904 | 1.947 | -2.77 | 3 |
| H413A | H10/11 | 28.1 | -0.8 | 0.454 | 0.088 | -1.31 | 3.294 | 0.381 | -2.59 | 1 |
| T427I | L-box | 26.2 | -2.7 | 0.037 | 0.018 | 2.31 | 3.307 | 0.561 | -2.58 | 8 |
| K421A-K422A | H10/11 | 26.4 | -2.5 | 0.037 | 0.009 | 2.31 | 3.346 | 0.264 | -2.57 | 6 |
| K421A | H10/11 | 24.9 | -4.0 | 0.043 | 0.004 | 2.09 | 4.412 | 1.176 | -2.17 | 3 |
| E435A | H12 | 28.9 | 0.0 | 0.340 | 0.026 | -0.89 | 4.628 | 0.960 | -2.10 | 2 |
| R314A | H6 | 26.5 | -2.4 | 0.083 | 0.031 | 1.14 | 5.084 | 1.219 | -1.96 | 5 |
| I420R | H10/11 | 28.0 | -0.9 | 0.655 | 0.105 | -1.84 | 5.205 | 1.049 | -1.93 | 1 |
| E329A | H7 | 27.2 | -1.7 | 0.207 | 0.038 | -0.18 | 6.277 | 0.847 | -1.66 | 1 |
| 72-441 | ΔAF1 | 27.0 | -1.9 | 0.379 | 0.105 | -1.05 | 6.586 | 3.113 | -1.59 | 3 |
| R147K | RFGR | 27.4 | -1.5 | 0.184 | 0.034 | -0.01 | 6.869 | 0.870 | -1.53 | 1 |
| T423A-T427D | L-box | 29.4 | 0.5 | 0.505 | 0.090 | -1.46 | 7.010 | 0.990 | -1.50 | 2 |
| R321K | between. H6/H7 | 28.1 | -0.8 | 0.780 | 0.023 | -2.09 | 7.312 | 2.103 | -1.44 | 3 |
| K421R | H10/11 | 26.9 | -2.0 | 0.095 | 0.005 | 0.95 | 7.632 | 1.816 | -1.38 | 2 |
| I327A | H7 | 31.6 | 2.7 | 0.675 | 0.201 | -1.88 | 7.950 | 3.227 | -1.32 | 3 |
| Y437F | H12 | 28.1 | -0.8 | 0.137 | 0.025 | 0.42 | 8.694 | 1.134 | -1.19 | 1 |
| T427F | L-box | 26.6 | -2.3 | 0.112 | 0.028 | 0.71 | 9.189 | 1.515 | -1.11 | 2 |
| E333A | H7 | 28.7 | -0.2 | 0.136 | 0.067 | 0.43 | 10.411 | 1.653 | -0.93 | 1 |
| L281A | H4 | 28.2 | -0.7 | 0.503 | 0.090 | -1.46 | 11.158 | 1.335 | -0.83 | 1 |
| K322R | between. H6/H7 | 28.4 | -0.5 | 0.186 | 0.038 | -0.02 | 12.295 | 1.734 | -0.69 | 1 |
| M417A | H10/11 | 29.4 | 0.5 | 0.515 | 0.083 | -1.49 | 12.405 | 2.419 | -0.68 | 2 |
| K421R-K422R | H10/11 | 27.4 | -1.5 | 0.104 | 0.025 | 0.82 | 13.107 | 0.726 | -0.60 | 2 |
| I420A | H10/11 | 29.5 | 0.6 | 0.354 | 0.118 | -0.95 | 13.269 | 2.499 | -0.58 | 1 |
| K422R | H10/11 | 28.0 | -0.9 | 0.182 | 0.066 | 0.01 | 13.455 | 0.093 | -0.56 | 2 |
| K318A | H6 | 28.8 | -0.1 | 0.199 | 0.044 | -0.12 | 13.830 | 2.605 | -0.52 | 1 |
| R314K | H6 | 28.8 | -0.1 | 0.163 | 0.033 | 0.17 | 14.221 | 1.954 | -0.48 | 1 |
| E424A | L-box | 28.8 | -0.1 | 0.262 | 0.084 | -0.52 | 15.137 | 3.025 | -0.39 | 1 |
| T427A | L-box | 28.2 | -0.7 | 0.140 | 0.038 | 0.39 | 15.137 | 1.517 | -0.39 | 1 |
| T427D | L-box | 31.5 | 2.6 | 0.465 | 0.138 | -1.35 | 15.263 | 5.367 | -0.38 | 5 |
| M416K | H10/11 | 28.0 | -0.9 | 0.268 | 0.040 | -0.55 | 15.455 | 1.913 | -0.36 | 1 |
| E426A | L-box | 27.9 | -1.0 | 0.306 | 0.065 | -0.74 | 15.560 | 1.320 | -0.35 | 1 |
| R154K | Hinge | 28.0 | -0.9 | 0.240 | 0.051 | -0.39 | 16.000 | 1.711 | -0.31 | 1 |
| E240A | H3 | 30.1 | 1.2 | 0.363 | 0.085 | -0.99 | 16.111 | 3.019 | -0.30 | 1 |
| E315A | H6 | 29.1 | 0.2 | 0.204 | 0.042 | -0.16 | 17.268 | 1.795 | -0.20 | 1 |
| T423D-T427D | L-box | 30.2 | 1.3 | 0.840 | 0.227 | -2.20 | 18.292 | 5.094 | -0.12 | 3 |
| K322A | between H6/H7 | 28.0 | -0.9 | 0.186 | 0.034 | -0.02 | 18.379 | 2.493 | -0.11 | 1 |
| K229R | between H2'/H3 | 28.5 | -0.4 | 0.291 | 0.034 | -0.67 | 19.027 | 1.684 | -0.06 | 1 |
| S428A | L-box | 27.4 | -1.5 | 0.156 | 0.051 | 0.23 | 19.160 | 4.785 | -0.05 | 1 |
| L432I | LXXLL, H12 | 30.3 | 1.4 | 0.525 | 0.229 | -1.52 | 19.562 | 6.226 | -0.02 | 1 |
| WT (Mean) | | 28.9 | 0.0 | 0.183 | 0.052 | 0.00 | 19.840 | 6.979 | 0.00 | 20 |
| K155R | Hinge | 28.4 | -0.5 | 0.193 | 0.098 | -0.08 | 20.393 | 2.208 | 0.04 | 1 |
| M416R | H10/11 | 30.5 | 1.6 | 0.218 | 0.056 | -0.25 | 22.943 | 4.186 | 0.21 | 1 |
| K300R | S2 | 29.0 | 0.1 | 0.149 | 0.083 | 0.30 | 23.918 | 2.663 | 0.27 | 1 |
| V254A | H3 | 30.4 | 1.5 | 0.651 | 0.225 | -1.83 | 24.062 | 7.498 | 0.28 | 2 |
| K300A | S2 | 29.1 | 0.2 | 0.295 | 0.094 | -0.69 | 24.930 | 2.444 | 0.33 | 1 |
| K318R | H6 | 29.2 | 0.3 | 0.124 | 0.018 | 0.56 | 25.634 | 4.667 | 0.37 | 1 |
| I328A | H7 | 30.7 | 1.8 | 0.406 | 0.028 | -1.15 | 26.355 | 1.582 | 0.41 | 1 |
| K239R | H3 | 29.6 | 0.7 | 0.283 | 0.084 | -0.63 | 28.247 | 3.346 | 0.51 | 1 |
| T423D-T427A | L-box | 28.8 | -0.1 | 0.230 | 0.039 | -0.33 | 28.791 | 0.989 | 0.54 | 3 |
| M416A | H10/11 | 28.9 | 0.0 | 0.358 | 0.070 | -0.97 | 30.484 | 2.966 | 0.62 | 1 |
| T423D | L-box | 29.2 | 0.3 | 0.257 | 0.042 | -0.49 | 30.484 | 4.059 | 0.62 | 1 |
| L433I | LXXLL, H12 | 30.9 | 2.0 | 0.742 | 0.054 | -2.02 | 31.651 | 24.418 | 0.67 | 3 |
| K422A | H10/11 | 28.7 | -0.2 | 0.133 | 0.013 | 0.46 | 32.728 | 4.286 | 0.72 | 2 |
| R321A | between H6/H7 | 30.8 | 1.9 | 0.547 | 0.061 | -1.58 | 33.591 | 3.374 | 0.76 | 1 |
| S428D | L-box | 29.3 | 0.4 | 0.342 | 0.054 | -0.90 | 36.002 | 5.205 | 0.86 | 1 |
| T425D | L-box | 30.1 | 1.2 | 0.374 | 0.075 | -1.03 | 41.355 | 5.207 | 1.06 | 1 |
| T423A | L-box | 31.1 | 2.2 | 0.678 | 0.049 | -1.89 | 43.518 | 9.927 | 1.13 | 2 |
| T425A | L-box | 29.9 | 1.0 | 0.426 | 0.060 | -1.22 | 44.632 | 5.241 | 1.17 | 1 |
| T423A-T427A | L-box | 29.8 | 0.9 | 0.415 | 0.172 | -1.18 | 48.961 | 18.688 | 1.30 | 2 |
| D326A | H7 | 30.3 | 1.4 | 0.454 | 0.092 | -1.31 | 49.180 | 7.973 | 1.31 | 1 |

| |
|---|
| wildtype |
| n > 1 |
| change ≤ -1,5 |
| change ≥ 1,5 |

Appendix D. Expression data for the PPARβ/δ mutant screen, sorted by effect on PT-S264–dependent change.

| mutant | position | basal repression | | PT-S264 | | | L165,041 | | | n |
|---|---|---|---|---|---|---|---|---|---|---|
| | | basal Ct | change vs WT | Mean | SD | change vs WT | MW | SD | change vs WT | |
| WT (Mean) | | 28.9 | 0.0 | 0.183 | 0.052 | 0.00 | 19.840 | 6.979 | 0.00 | 20 |
| L406R | H10/11 | 26.5 | -2.4 | 1.057 | 0.111 | -2.53 | 1.206 | 0.124 | -4.04 | 1 |
| C91A-E92A | DBD | 27.3 | -1.6 | 1.020 | 0.194 | -2.48 | 1.220 | 0.173 | -4.02 | 8 |
| L432A | LXXLL, H12 | 30.8 | 1.9 | 1.020 | 0.392 | -2.48 | 2.904 | 0.147 | -2.77 | 3 |
| V257R | H3 | 27.0 | -1.9 | 1.012 | 0.142 | -2.47 | 1.250 | 0.260 | -3.99 | 3 |
| empty vector | | 26.3 | -2.6 | 0.993 | 0.116 | -2.44 | 1.273 | 0.234 | -3.96 | 9 |
| T261R | H3 | 26.8 | -2.1 | 0.982 | 0.136 | -2.42 | 1.366 | 0.179 | -3.86 | 3 |
| A404K | H10/11 | 27.1 | -1.8 | 0.979 | 0.104 | -2.42 | 2.346 | 0.263 | -3.08 | 1 |
| L429A | LXXLL | 29.9 | 1.0 | 0.926 | 0.424 | -2.34 | 2.904 | 1.947 | -2.77 | 3 |
| H413W | H10/11 | 28.5 | -0.4 | 0.920 | 0.090 | -2.33 | 1.196 | 0.189 | -4.05 | 5 |
| T261R-V279R | H3/H4 | 26.7 | -2.2 | 0.898 | 0.016 | -2.29 | 1.035 | 0.007 | -4.26 | 2 |
| G146A | RFGR | 26.0 | -2.9 | 0.877 | 0.090 | -2.26 | 1.266 | 0.060 | -3.97 | 1 |
| T423D-T427D | L-box | 30.2 | 1.3 | 0.840 | 0.227 | -2.20 | 18.292 | 5.094 | -0.12 | 3 |
| D439A | H12 | 26.3 | -2.6 | 0.836 | 0.120 | -2.19 | 1.486 | 0.360 | -3.74 | 3 |
| R321K | between. H6/H7 | 28.1 | -0.8 | 0.780 | 0.023 | -2.09 | 7.312 | 2.103 | -1.44 | 3 |
| A404W | H10/11 | 27.4 | -1.5 | 0.774 | 0.086 | -2.08 | 2.549 | 0.252 | -2.96 | 1 |
| L433I | LXXLL, H12 | 30.9 | 2.0 | 0.742 | 0.054 | -2.02 | 31.651 | 24.418 | 0.67 | 3 |
| L433F | LXXLL, H12 | 31.0 | 2.1 | 0.708 | 0.164 | -1.95 | 1.382 | 0.204 | -3.84 | 3 |
| T423A | L-box | 31.1 | 2.2 | 0.678 | 0.049 | -1.89 | 43.518 | 9.927 | 1.13 | 2 |
| I327A | H7 | 31.6 | 2.7 | 0.675 | 0.201 | -1.88 | 7.950 | 3.227 | -1.32 | 3 |
| I420R | H10/11 | 28.0 | -0.9 | 0.655 | 0.105 | -1.84 | 5.205 | 1.049 | -1.93 | 1 |
| V254A | H3 | 30.4 | 1.5 | 0.651 | 0.225 | -1.83 | 24.062 | 7.498 | 0.28 | 2 |
| V279R | H4 | 26.8 | -2.1 | 0.650 | 0.030 | -1.83 | 1.128 | 0.077 | -4.14 | 3 |
| Y437A | H12 | 29.5 | 0.6 | 0.591 | 0.050 | -1.69 | 1.461 | 0.234 | -3.76 | 3 |
| R321A | between H6/H7 | 30.8 | 1.9 | 0.547 | 0.061 | -1.58 | 33.591 | 3.374 | 0.76 | 1 |
| F324A | between H6/H7 | 28.1 | -0.8 | 0.540 | 0.183 | -1.56 | 2.269 | 0.198 | -3.13 | 3 |
| L433A | LXXLL, H12 | 29.4 | 0.5 | 0.532 | 0.161 | -1.54 | 1.283 | 0.287 | -3.95 | 1 |
| L432I | LXXLL, H12 | 30.3 | 1.4 | 0.525 | 0.229 | -1.52 | 19.562 | 6.226 | -0.02 | 1 |
| K283A | H4 | 28.7 | -0.2 | 0.516 | 0.059 | -1.50 | 2.782 | 0.823 | -2.83 | 2 |
| M417A | H10/11 | 29.4 | 0.5 | 0.515 | 0.083 | -1.49 | 12.405 | 2.419 | -0.68 | 2 |
| T423A-T427D | L-box | 29.4 | 0.5 | 0.505 | 0.090 | -1.46 | 7.010 | 0.990 | -1.50 | 2 |
| L281A | H4 | 28.2 | -0.7 | 0.503 | 0.090 | -1.46 | 11.158 | 1.335 | -0.83 | 1 |
| T427D | L-box | 31.5 | 2.6 | 0.465 | 0.138 | -1.35 | 15.263 | 5.367 | -0.38 | 5 |
| V254R | H3 | 29.6 | 0.7 | 0.464 | 0.106 | -1.34 | 1.508 | 0.151 | -3.72 | 2 |
| H413A | H10/11 | 28.1 | -0.8 | 0.454 | 0.088 | -1.31 | 3.294 | 0.381 | -2.59 | 1 |
| D326A | H7 | 30.3 | 1.4 | 0.454 | 0.092 | -1.31 | 49.180 | 7.973 | 1.31 | 1 |
| 1-428 | ΔAF2 | 28.5 | -0.4 | 0.426 | 0.079 | -1.22 | 1.040 | 0.199 | -4.25 | 3 |
| T425A | L-box | 29.9 | 1.0 | 0.426 | 0.060 | -1.22 | 44.632 | 5.241 | 1.17 | 1 |
| T423A-T427A | L-box | 29.8 | 0.9 | 0.415 | 0.172 | -1.18 | 48.961 | 18.688 | 1.30 | 2 |
| I328A | H7 | 30.7 | 1.8 | 0.406 | 0.028 | -1.15 | 26.355 | 1.582 | 0.41 | 1 |
| 1-421 | ΔL-box+AF2 | 28.6 | -0.3 | 0.382 | 0.044 | -1.06 | 1.233 | 0.068 | -4.01 | 3 |
| 72-441 | ΔAF1 | 27.0 | -1.9 | 0.379 | 0.105 | -1.05 | 6.586 | 3.113 | -1.59 | 3 |
| T425D | L-box | 30.1 | 1.2 | 0.374 | 0.075 | -1.03 | 41.355 | 5.207 | 1.06 | 1 |
| E240A | H3 | 30.1 | 1.2 | 0.363 | 0.085 | -0.99 | 16.111 | 3.019 | -0.30 | 1 |
| M416A | H10/11 | 28.9 | 0.0 | 0.358 | 0.070 | -0.97 | 30.484 | 2.966 | 0.62 | 1 |
| I420A | H10/11 | 29.5 | 0.6 | 0.354 | 0.118 | -0.95 | 13.269 | 2.499 | -0.58 | 1 |
| S428D | L-box | 29.3 | 0.4 | 0.342 | 0.054 | -0.90 | 36.002 | 5.205 | 0.86 | 1 |
| E435A | H12 | 28.9 | 0.0 | 0.340 | 0.026 | -0.89 | 4.628 | 0.960 | -2.10 | 2 |
| E426A | L-box | 27.9 | -1.0 | 0.306 | 0.065 | -0.74 | 15.560 | 1.320 | -0.35 | 1 |
| K300A | S2 | 29.1 | 0.2 | 0.295 | 0.094 | -0.69 | 24.930 | 2.444 | 0.33 | 1 |
| K229R | between H2'/H3 | 28.5 | -0.4 | 0.291 | 0.034 | -0.67 | 19.027 | 1.684 | -0.06 | 1 |
| K239R | H3 | 29.6 | 0.7 | 0.283 | 0.084 | -0.63 | 28.247 | 3.346 | 0.51 | 1 |
| H430K | LXXLL | 30.1 | 1.2 | 0.271 | 0.126 | -0.57 | 2.770 | 0.444 | -2.84 | 1 |
| M416K | H10/11 | 28.0 | -0.9 | 0.268 | 0.040 | -0.55 | 15.455 | 1.913 | -0.36 | 1 |
| E424A | L-box | 28.8 | -0.1 | 0.262 | 0.084 | -0.52 | 15.137 | 3.025 | -0.39 | 1 |
| T423D | L-box | 29.2 | 0.3 | 0.257 | 0.042 | -0.49 | 30.484 | 4.059 | 0.62 | 1 |
| R154K | Hinge | 28.0 | -0.9 | 0.240 | 0.051 | -0.39 | 16.000 | 1.711 | -0.31 | 1 |
| T423D-T427A | L-box | 28.8 | -0.1 | 0.230 | 0.039 | -0.33 | 28.791 | 0.989 | 0.54 | 3 |
| M416R | H10/11 | 30.5 | 1.6 | 0.218 | 0.056 | -0.25 | 22.943 | 4.186 | 0.21 | 1 |
| E329A | H7 | 27.2 | -1.7 | 0.207 | 0.038 | -0.18 | 6.277 | 0.847 | -1.66 | 1 |
| E315A | H6 | 29.1 | 0.2 | 0.204 | 0.042 | -0.16 | 17.268 | 1.795 | -0.20 | 1 |
| K318A | H6 | 28.8 | -0.1 | 0.199 | 0.044 | -0.12 | 13.830 | 2.605 | -0.52 | 1 |
| K155R | Hinge | 28.4 | -0.5 | 0.193 | 0.098 | -0.08 | 20.393 | 2.208 | 0.04 | 1 |
| K322A | between H6/H7 | 28.0 | -0.9 | 0.186 | 0.034 | -0.02 | 18.379 | 2.493 | -0.11 | 1 |
| K322R | between. H6/H7 | 28.4 | -0.5 | 0.186 | 0.038 | -0.02 | 12.295 | 1.734 | -0.69 | 1 |
| R147K | RFGR | 27.4 | -1.5 | 0.184 | 0.034 | -0.01 | 6.869 | 0.870 | -1.53 | 1 |
| WT (Mean) | | 28.9 | 0.0 | 0.183 | 0.052 | 0.00 | 19.840 | 6.979 | 0.00 | 20 |
| K422R | H10/11 | 28.0 | -0.9 | 0.182 | 0.066 | 0.01 | 13.455 | 0.093 | -0.56 | 2 |
| R314K | H6 | 28.8 | -0.1 | 0.163 | 0.033 | 0.17 | 14.221 | 1.954 | -0.48 | 1 |
| S428A | L-box | 27.4 | -1.5 | 0.156 | 0.051 | 0.23 | 19.160 | 4.785 | -0.05 | 1 |
| K300R | S2 | 29.0 | 0.1 | 0.149 | 0.083 | 0.30 | 23.918 | 2.663 | 0.27 | 1 |
| T427A | L-box | 28.2 | -0.7 | 0.140 | 0.038 | 0.39 | 15.137 | 1.517 | -0.39 | 1 |
| Y437F | H12 | 28.1 | -0.8 | 0.137 | 0.025 | 0.42 | 8.694 | 1.134 | -1.19 | 1 |
| E333A | H7 | 28.7 | -0.2 | 0.136 | 0.067 | 0.43 | 10.411 | 1.653 | -0.93 | 1 |
| K422A | H10/11 | 28.7 | -0.2 | 0.133 | 0.013 | 0.46 | 32.728 | 4.286 | 0.72 | 2 |
| K318R | H6 | 29.2 | 0.3 | 0.124 | 0.018 | 0.56 | 25.634 | 4.667 | 0.37 | 1 |
| T427F | L-box | 26.6 | -2.3 | 0.112 | 0.028 | 0.71 | 9.189 | 1.515 | -1.11 | 2 |
| K421R-K422R | H10/11 | 27.4 | -1.5 | 0.104 | 0.025 | 0.82 | 13.107 | 0.726 | -0.60 | 2 |
| K421R | H10/11 | 26.9 | -2.0 | 0.095 | 0.005 | 0.95 | 7.632 | 1.816 | -1.38 | 2 |
| R314A | H6 | 26.5 | -2.4 | 0.083 | 0.031 | 1.14 | 5.084 | 1.219 | -1.96 | 5 |
| K421A | H10/11 | 24.9 | -4.0 | 0.043 | 0.004 | 2.09 | 4.412 | 1.176 | -2.17 | 3 |
| T427I | L-box | 26.2 | -2.7 | 0.037 | 0.018 | 2.31 | 3.307 | 0.561 | -2.58 | 8 |
| K421A-K422A | H10/11 | 26.4 | -2.5 | 0.037 | 0.009 | 2.31 | 3.346 | 0.264 | -2.57 | 6 |

- wildtype
- n > 1
- change ≤ -1,5
- change ≥ 1,5
